# Profiling The Compendium Of Changes In *Saccharomyces cerevisiae* Due To Mutations That Alter Availability Of The Main Methyl Donor S-Adenosylmethionine

**DOI:** 10.1101/2023.06.09.544294

**Authors:** McKayla Remines, Makailyn Schoonover, Zoey Knox, Kailee Kenwright, Kellyn M. Hoffert, Amila Coric, James Mead, Joseph Ampfer, Serigne Seye, Erin D. Strome

**Affiliations:** Department of Biological Sciences, Northern Kentucky University, Highland Heights, KY 41099

## Abstract

The *SAM1* and *SAM2* genes encode for S-AdenosylMethionine (AdoMet) synthetase enzymes, with AdoMet serving as the main methyl donor. We have previously shown that independent deletion of these genes alters chromosome stability and AdoMet concentrations in opposite ways in *S. cerevisiae.* To characterize other changes occurring in these mutants, we grew wildtype, *sam1*Δ*/sam1*Δ, and *sam2*Δ*/sam2*Δ strains in 15 different Phenotypic Microarray plates with different components, equal to 1440 wells, and measured for growth variations. RNA-Sequencing was also carried out on these strains and differential gene expression determined for each mutant. In this study, we explore how the phenotypic growth differences are linked to the altered gene expression, and thereby predict the mechanisms by which loss of the *SAM* genes and subsequent AdoMet level changes, impact *S. cerevisiae* pathways and processes. We present six stories, discussing changes in sensitivity or resistance to azoles, cisplatin, oxidative stress, arginine biosynthesis perturbations, DNA synthesis inhibitors, and tamoxifen, to demonstrate the power of this novel methodology to broadly profile changes due to gene mutations. The large number of conditions that result in altered growth, as well as the large number of differentially expressed genes with wide-ranging functionality, speaks to the broad array of impacts that altering methyl donor abundance can impart, even when the conditions tested were not specifically selected as targeting known methyl involving pathways. Our findings demonstrate that some cellular changes are directly related to AdoMet-dependent methyltransferases and AdoMet availability, some are directly linked to the methyl cycle and its role is production of several important cellular components, and others reveal impacts of *SAM* gene mutations on previously unconnected pathways.

**AUTHOR SUMMARY:** S-AdenosylMethionine, or AdoMet, is the main methyl donor in all cells. Methylation reactions are used broadly and impact numerous processes and pathways. The *SAM1* and *SAM2* genes of *Saccharomyces cerevisiae* are responsible for producing the enzymes called S-Adenosylmethionine synthetases, which make AdoMet from methionine and ATP. Our previous research showed that when these genes are deleted independently, they have opposite effects on AdoMet levels and chromosome stability. To advance our understanding of the wide array of changes going on in cells with these gene deletions we characterized our mutants phenotypically, growing in various different conditions to look for growth changes, and for their different gene expression profiles. In this study, we investigated how the differences in growth patterns are connected to the altered gene expression, and thereby were able to predict the mechanisms through which the loss of the *SAM* genes affects different pathways. Our investigations have uncovered novel mechanisms of sensitivity or resistance to many conditions and shown linkages to AdoMet availability, AdoMet-dependent methyltransferases, methyl cycle compounds, or new connections to *sam1* and *sam2* gene deletions.

## INTRODUCTION

*SAM1* and *SAM2* are paralogous genes in *Saccharomyces cerevisiae*, which encode proteins responsible for the synthesis of S-AdenosylMethionine (AdoMet), the major methyl donor in all cells (Fig 1A). Cells with either *sam1* or *sam2* deletions are able to survive, however, the double homozygous deletion of *sam1* and *sam2* renders cells inviable [1]. *SAM* gene mutations have been identified to alter AdoMet abundance; *sam1*-deficient cells have been found to have increased AdoMet levels, whereas *sam2*-deficient cells have decreased AdoMet levels [2]. Methylation events, using methyl donors such as AdoMet, are used to regulate many cellular processes. In *S. cerevisiae,* the cellular targets regulated by methylation events include RNAs, proteins, lipids, nucleotides and small molecules [3] (Fig 1B). Methylation requires a methyltransferase enzyme, and many AdoMet-dependent methyltransferases exist in *S. cerevisiae* which carry out a wide variety and number of these reactions.

**Fig 1.**
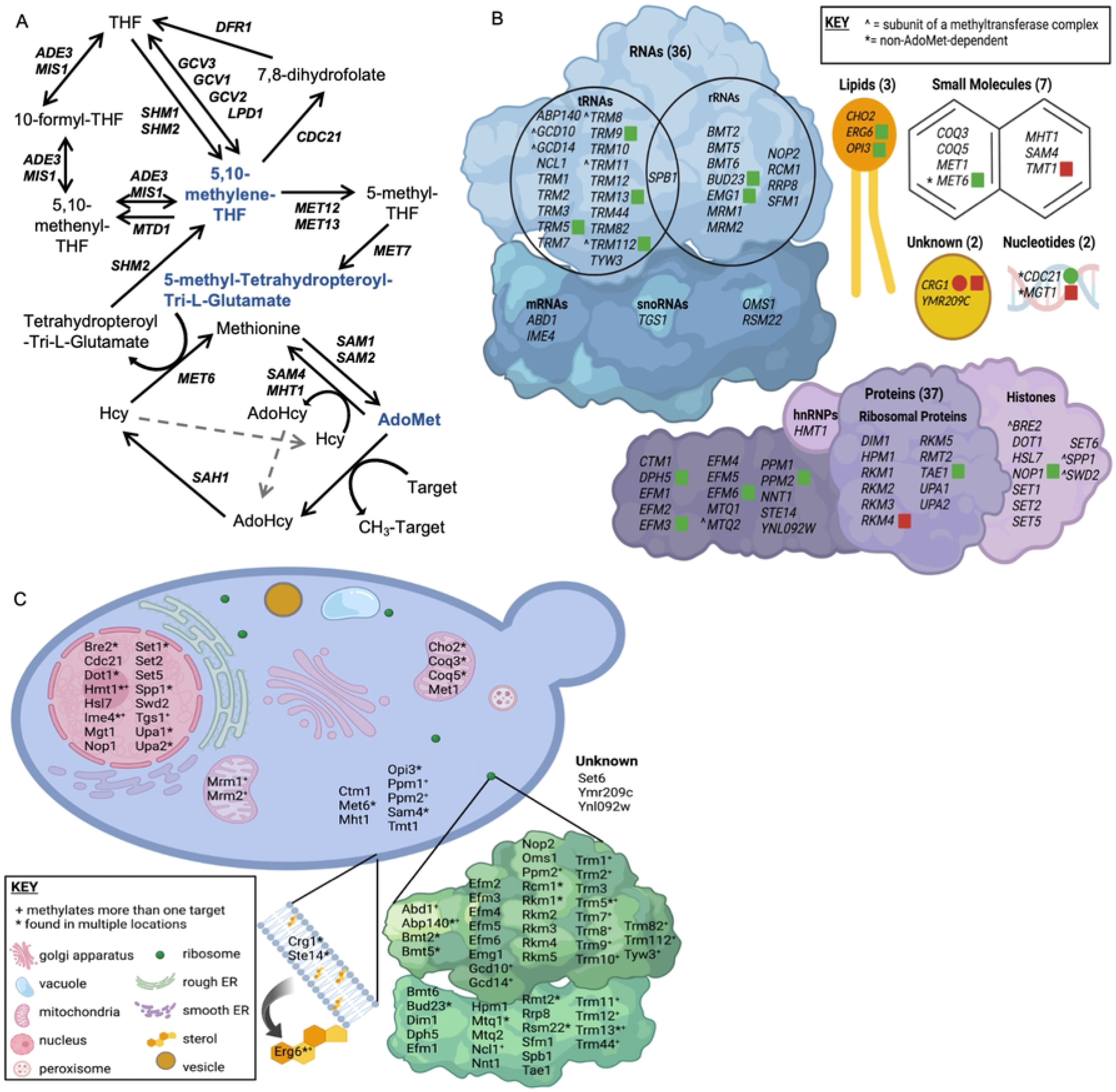
Methyl donors, me+thyltransferases, and methylation target molecules of *S. cerevisiae*. (A) The methyl cycle in *S. cerevisiae* which produces the main methyl donor AdoMet, and portions of the folate cycle that produce the less frequently used methyl donors, 5,10-methylenetetrahydrofolate and 5-methyltetrahydrofolate. Methyl donors are in blue. (B) There are 87 identified methyltransferases in budding yeast, with the target macromolecule identified for most. The main targets are proteins and RNAs, followed by small molecules, lipids, and nucleotides, with 2 methyltransferases with unconfirmed targets. (C) The methyltransferases are found in a wide variety of locations in the cell. The “*” next to methyltransferase names identify enzymes that are active in more than one location/organelle. The “+” identifies enzymes that methylate multiple target molecules. B & C Created with BioRender.com

There are 87 identified methyltransferases in *S. cerevisiae,* all but three are AdoMet-dependent, and 84 contain known or putative substrates (Fig 1B). Of the AdoMet-dependent methyltransferases, 42.9% target RNA molecules, 44% target proteins, 3.6% have lipid targets, 7.1% target small molecules, and 2.4% remain with their target unidentified. Methyltransferases play roles in a wide variety of processes in the cell including translation and ribosome synthesis, RNA processing, DNA maintenance, amino acid metabolism, respiration, vesicle transport, autophagy, lipid homeostasis, sterol synthesis, and sulfur and nitrogen reduction (Fig 1C). This wide range of targets, along with the large variety of impacts of the methylation events themselves, lead to broad effects in the cell and are not limited to impacting only a few pathways or cellular outcomes.

Among the AdoMet-dependent RNA methyltransferases, 19 target tRNAs, one targets both tRNAs and rRNAs, 11 target rRNAs, two target mRNAs, two are predicted to target RNA, and one targets sn/snoRNAs [4–37]. When methylation is targeted to tRNAs it has been shown to function in folding dynamics, thermostability, structure stabilization, protection from cleavage, maturation, regulation of degradation and localization, charging fidelity, and codon-anticodon interactions [38,39]. When targeted to rRNAs, methylation functions in protection from hydrolytic cleavage, structure stabilization, helix flexibility, accessibility, and regulation of a subset of mRNAs [40]. Methylation of mRNA can occur on the 5’ cap and functions in termination of transcription, efficient mRNA splicing, and recruitment of protein complexes required for exosome degradation, translation, and ultimately cell growth [4,41]. Other mRNA methylation is restricted to meiotically dividing yeast and has been shown to function in the induction of meiosis and the suppression of pseudohyphal growth under conditions of nutrient deprivation [42]. As well, in meiosis, mRNA methylation is found most often on genes related to DNA replication, mismatch repair, and synaptonemal complex formation but is also found on genes related to signaling, maintenance, and metabolism [43]. The transcripts that are enriched in methylation have been determined to be the most efficiently translated [44]. Hypermethylation of the m^7^G cap structure to m_3_G on U1, U2, U4, and U5 snRNAs as well as U3 and snR4 of the C/D class of snoRNAs and snR8, snR11, snR31, snR33, snR35, and snR42 of the H/ACA class of snoRNAs have been shown to be required at cold temperatures for normal localization and splicing processes of mitotically dividing yeast [11]. More importantly, the m_3_G cap on these sn/snoRNAs are required for implementation of meiosis and sporulation via promoting efficient splicing of meiotic pre-mRNAs [45].

Of the 37 AdoMet-dependent protein methyltransferases, the majority have been found to target ribosomal proteins, however other targets include histone proteins, translation factors such as elongation and release factors, cytochrome c, heterogeneous nuclear ribonucleoproteins (hnRNPs), phosphoprotein phosphatase 2A (PP2A), α-factor mating pheromone, RAS proteins, as well as a subunit of the DASH complex of kinetochores [30,34,36,46–74]. Methylation of histone proteins has been shown to function in gene regulation, regulation of chromatin silencing, regulation of translesion synthesis to maintain genome integrity, nucleosome formation, cell-cycle checkpoints, repression of bidirectional transcription and aberrant RNA polymerase II localization to prevent cryptic transcripts, and DNA repair occurring at double-strand breaks, uncapped telomeres, and in response to ultraviolet induced damage [47,49,50,75]. When methylation is targeted to ribosomal proteins, it may play a role in regulating interactions between proteins and rRNA and/or tRNA, protecting cells during translational stress, ribosome biogenesis, structural stabilization, folding conformations and dynamics, as well as accuracy of translation. Much is still unknown about the function of methylation of translation factors, but it may play a role in viral replication pathways, structural stabilization, as well as translational function and accuracy. Methylation of cytochrome c has been shown to regulate its function via promoting association with other proteins as well as facilitating its import into the mitochondria [46,48]. hnRNP methylation aids in its export from the nucleus [48,51]. Methylation of PP2A is required for complex formation and therefore functionality [52,53]. Methylation of α-factor mating pheromone and RAS proteins may play a role in maturation efficiency and localization [54]. Lastly, methylation of the protein subunit of the DASH complex has been shown to be important for regulation of kinetochore-microtubule attachments that lead to proper chromosomal segregation [46,47].

Three AdoMet-dependent lipid methyltransferases are known to exist, and lipid methylation has been implicated in the conversion of zymosterol to fecosterol within the ergosterol biosynthesis pathway which is necessary for accurate endocytosis, cell polarization, cell fusion, and cell wall assembly [76–78]. Lipid methylation has also been implicated in phospholipid biosynthesis during the conversion of phosphatidylethanolamine to phosphatidylcholine which has been linked to induction of mitophagy [78,79]. Ordered mitophagy is essential for mitochondrial quantity and quality control to maintain energy requirements for the cell as well as protection from reactive oxygen species (ROS) and damaged organelles [76,79]. Additionally, phosphatidylcholine is an important phospholipid in membranes; it provides structure and causes curvature that is used in budding and fusion [80]. Phosphatidylcholine has also been shown to be required for the functionality of the sorting and assembly machinery complex in the mitochondria [81].

Six AdoMet-dependent small molecule methyltransferases have been identified with small molecule methylation functioning in the biosynthesis of ubiquinone (CoEnzyme Q) and siroheme, as well as in the detoxification of the citric acid cycle, the amino acid stress response leading to invasive growth, and the control of methionine/AdoMet ratios. One methyltransferase involved in methylation of small molecule homocysteine, *MET6,* is AdoMet-independent. In the absence of methionine synthase encoded by *MET6* or in the presence of extracellular AdoMet, Sam4 and Mht1 use AdoMet as a methyl donor to synthesize methionine from homocysteine. AdoMet/methionine ratios are important for overall cellular homeostasis due to the extensive roles AdoMet plays in the cell as well as the importance of methionine in one-carbon metabolism. Siroheme is necessary for sulfite reductase activity to assimilate sulfur and subsequently synthesize sulfur amino acids. Defects in siroheme biosynthesis, including disrupted methyltransferase activity of Met1, can lead to cell death, as siroheme is required for growth. Lack of gene products from the *COQ5* and *COQ3* AdoMet-dependent methyltransferases lead to cellular deficiency in ubiquinone and arrest of aerobic respiration [82–88]. *TMT1* encodes for a trans-aconitate methyltransferase enzyme, which catalyzes the methyl esterification of trans-aconitate and 3-isopropylmate. Trans-aconitate is a known inhibitor of the citric acid cycle, and it is believed that methylation of this molecule results in decreased inhibition on the citric acid cycle [83]. 3-isopropylmate is an intermediate in the leucine biosynthetic pathway, though the biological significance of the methylation of this molecule has yet to be discovered [82].

While AdoMet is the main methyl donor, there are three methyltransferases that are not AdoMet-dependent: Cdc21, Met6, and Mgt1. Cdc21 and Mgt1 are both methyltransferases that are responsible for methylation of nucleotides. Cdc21 synthesizes dTMP from dUMP using 5,10-methylenetetrahydrofolate (5,10-CH_2_-THF) as the methyl donor, while Met6 synthesizes methionine from homocysteine using polyglutamated versions of 5-methyltetrahydrofolate as the methyl donor [89,90]. Mgt1 is responsible for methyltransferase activity in DNA damage repair and protection against alkylation damage, and accepts methyl groups from 6-O-methylguanine to form S-methylcysteine [74,91]. Interestingly, these non-AdoMet-dependent methyltransferases may still be impacted by changes in AdoMet. For example, the methyl donors for Cdc21 and Met6 are produced within the folate cycle which stems directly from the methyl cycle (Fig 1). Therefore, alterations in AdoMet abundance could impact the abundance of these alternate methyl donor compounds as well. As previously mentioned, the small molecule methyltransferase Mht1 can use AdoMet for its function, however it shows a strong preference for *S*-methylmethionine (SMM). SMM has only been shown to be produced in plants, however Thomas et al. theorize that yeast cells living on plants evolved Mht1, and Mmp1 the SMM permease, as a mechanism to use plant-derived SMM [84].

The vast majority of the AdoMet-dependent methyltransferases are regulated at the transcriptional level via transcriptional activators and repressors. Several are regulated via chromatin modification such as acetylation, deacetylation, and interaction with silencer proteins [73,92–94]. Coq5, a small molecule methyltransferase, is regulated at the post-translational level via cleavage of a destabilizing N-terminal residue upon import to the mitochondria [95]. Trm4, a tRNA methyltransferase, is also regulated via protein modification through phosphorylation of a threonine residue [96]. While SAM/AdoMet riboswitches have been studied intensively in bacteria, similar riboswitches have not been found in *S. cerevisiae* [97]. Further, while compounds of the methyl cycle have been shown to be involved in transcriptional responses, the exact mechanisms of their involvement have not yet been elucidated, and it is unclear what role AdoMet might directly play in the control mechanisms for production or activation of the AdoMet-dependent methyltransferases [98].

Beyond its use as a methyl donor, AdoMet has proven to play a key role in other cellular and metabolic pathways, including glutathione (GSH), polyamine, and dNTP synthesis. AdoMet is produced from Methionine (Met) and ATP. When an AdoMet-dependent methyltransferase uses AdoMet as a methyl donor and transfers off the methyl group, the byproduct is S-AdenosylHomocysteine (AdoHcy) (Fig 1A). AdoHcy hydrolase catabolizes AdoHcy to form homocysteine (Hcy) which can then be converted back to methionine by Met6 using a methyl donor from the folate cycle. Hcy can also be used to synthesize several metabolic intermediates vital to cellular survival, so there is a balance that must be achieved between conversion back into methionine or use in transsulfuration. The transsulfuration pathway is responsible for production of GSH from Hcy, producing cysteine and important signaling molecules as intermediates [99]. GSH assists in protection from ROS within the cell, detoxification, damage repair, and enzyme production [100]. ROS is a common byproduct of mitochondrial oxidative metabolism during ATP production and can lead to cellular damage or death after prolonged exposure. Homocysteine conversion back to methionine involves products from the folate cycle, specifically a methylated, poly-glutamated, tetrahydrofolate (THF). THF production and conversion amongst derivative forms comprises the folate cycle (Fig 1A). THF, an amino acid derivative, is an important molecule in synthesis of amino acids, nucleic acids, and nitrogenous bases, including pyrimidines and purines [101]. A THF derivative, 5-methyltetrahydropteroyl-tri-L-glutamate, appears to be the most commonly used methyl donor when Hcy is converted to Met by Met6. Met6 requires a methylated THF derivative with at least two glutamates attached [90]. The now unmethylated poly-glutamate THF derivative can then be recycled back into the folate cycle through use of Shm2. The folate cycle also supplies 5,10-methylene-THF to be used as the methyl donor for Cdc21 conversion of dUMP to dTMP, and thus plays a vital role in nucleotide synthesis. Altered dNTP levels have been shown to cause a variety of problems that results in increased damage to DNA and chromosomal instability. Aminopropylation metabolizes AdoMet to synthesize polyamines necessary for cellular functions important for cell growth [102]. The most common polyamines are putrescine, spermidine, and spermine, each consisting of a core molecule with three or more amino groups attached, and carrying a positive charge [103]. Putrescine is synthesized either directly from ornithine, by ornithine decarboxylase, or indirectly from arginine, by arginine decarboxylase. For additional polyamine synthesis, AdoMet is then decarboxylated and the aminopropyl group is transferred to the putrescine molecule to produce spermidine. An additional AdoMet molecule can then be used for another aminopropyl group transfer to spermidine to form spermine. During polyamine synthesis, a by-product called methylthioadenosine is produced. Methylthioadenosine can be converted back to methionine by a series of enzymes within the methionine salvage pathway. Spermidine and spermine are crucial for stabilizing DNA and RNA [104]. Stabilization of DNA occurs via binding to the minor groove of nucleic acids, driven by electrostatic interaction, especially if the DNA is B-form or under stress [105]. It is thought that polyamines play a role in gene regulation through their interaction with the backbone of DNA, which in turn effects how well DNA-binding proteins can target their binding sites [106]. Yet another vital biological process involving polyamines is that of membrane stabilization. Polyamines, especially spermine molecules, can stabilize the plasma membrane, particularly during osmotic salt stress. It is hypothesized that they do so by maintaining, and even improving, ionic equilibrium through modification of the plasma membrane, including adjusting calcium ion channel allocation to halt entry of sodium and potassium ions into the cytoplasm [107]. It has further been identified that polyamines stabilize the plasma membrane by decreasing deformability and providing resistance to fragmentation [108]. Other cellular functions that polyamines participate in include roles in signal transduction, stress response and homeostasis, RNA synthesis, and protein interactions [107].

Our group has previously found that AdoMet concentration perturbances, due to the independent loss of *sam1* or *sam2*, result in changes to the stability of the genome as measured with chromosome stability assays [2]. Interestingly, *sam1* deletion results in increased AdoMet concentrations and a more stable genome while the removal of *sam2* decreased AdoMet concentrations and resulted in a less stable genome. Due to the wide array of processes in the cell that are impacted by AdoMet availability, including the aforementioned methylation events, as well as the key cellular molecules produced from products of the methyl cycle, we sought to broadly characterize the multitude of changes occurring in cells due to the *sam1* and *sam2* gene deletions. To accomplish this, we have undertaken phenotypic profiling and gene expression analysis in these mutant strains. Phenotypic profiling involved the Phenotypic Microarray system from BiOLOG, and characterizing growth differences of mutant and wildtype strains in over 1,000 different conditions [109]. Observed growth differences were then explored to understand the mechanism of action of the causative condition and its relation to methylation events, methyl cycle products, and/or AdoMet availability. We also undertook RNA-Sequencing experiments to understand the impacts of AdoMet availability on gene expression. As described above, many methylation events can impact gene expression, including via histone methylation and mRNA methylation. Therefore, exploration of differential gene expression in our mutant strains also allows us to characterize impacted genes/pathways with no intrinsic relationship to methylation, the methyl cycle, or AdoMet beyond their regulation. Further, we sought to link these two methodologies, aiming to align these data sets to elucidate where gene expression differences explain altered phenotypes. Through the combination of these methodologies, we have built a more comprehensive understanding of the changes occurring due to the *sam1* and *sam2* deletions and are able to hypothesize which of these altered pathways might be contributing to the observed impacts on genome stability.

## RESULTS

### RNA-Sequencing

RNA-Sequencing revealed 134 genes in the *sam1Δ/sam1Δ* mutant strain whose expression patterns differed from wildtype baseline expression. Specifically, there were 59 upregulated genes and 75 downregulated genes at the following cutoff values: p-value ≤ 0.05, 0.67 ≤ |Fold Change (FC)| ≥ 1.5. A list and description of all upregulated and downregulated genes can be found in Supplementary Table 1. Gene Ontology analysis of the list from the *sam1Δ/sam1Δ* strain revealed that the differentially expressed genes were enriched for several terms across the three domains (Table 1). When looked at altogether, it appears that the deletion of both copies of *sam1* leads to differential expression of genes that might play roles in protein and xenobiotic metabolism as well as transmembrane transport of ions. Only one AdoMet-dependent methyltransferase gene, *CRG1*, was found to be altered in its expression between the *sam1*-deficient strain and wildtype, being upregulated 1.58-fold (p-value = 0.0236) (Fig 1B). While the substrate of this protein is still unknown, Crg1 is known to be involved in lipid homeostasis [110].

**Table 1.** GOConsortium Gene Ontology categorization of differentially expressed genes from RNA-sequencing. GOConsortium was used to analyze *sam1Δ/sam1Δ* and *sam2Δ/sam2Δ* DEGs for enriched and underrepresented biological process, molecular function, and cellular component gene ontology terms at a p-value and FDR cutoff of 0.05 [111]. Overrepresented terms are indicated with “+” whereas underrepresented terms are indicated with “-”. The current table lists the most specific subclass of enriched terms for each category. A complete list of enriched terms, including parent classes, can be found in Supplementary Table 2.

| Strain | GO Category | GO: ID | Description | +/- | p-value | FDR |
| --- | --- | --- | --- | --- | --- | --- |
| *sam1Δ/sam1Δ* | Biological Process | GO:0006805 | Xenobiotic metabolic process | + | 1.23e-04 | 3.56e-02 |
|  |  | GO:0044718 | Siderophore transmembrane transport | + | 1.83e-04 | 4.76e-02 |
|  |  | GO:0009063 | Cellular amino acid catabolic process | + | 1.34e-04 | 3.67e-02 |
|  |  | GO:0044267 | Cellular protein metabolic process | + | 8.91e-05 | 2.73e-02 |
|  |  | GO:0010467 | Gene expression | + | 7.39e-06 | 9.63e-03 |
|  | Molecular Function | GO:0015293 | Symporter activity | + | 4.73e-05 | 3.00e-02 |
|  |  | GO:0015075 | Ion transmembrane transporter activity | + | 4.46e-05 | 3.77e-02 |
|  | Cellular Component | GO:0000786 | Nucleosome | + | 4.65e-04 | 4.95e-02 |
|  |  | GO:0005887 | Integral component of plasma membrane | + | 4.34-e06 | 4.61e-03 |
|  |  | GO:0000324 | Fungal-type vacuole | + | 4.51e-05 | 1.60e-02 |
| **sam2Δ/sam2Δ** | Biological Process | GO:0006526 | Arginine biosynthetic process | + | 2.89e-05 | 5.57e-03 |
|  |  | GO:0006591 | Ornithine biosynthetic process | + | 8.69e-05 | 1.37e-02 |
|  |  | GO:0000097 | Sulfur amino acid biosynthetic process | + | 1.09e-04 | 1.58e-02 |
|  |  | GO:0072528 | Pyrimidine-containing compound biosynthetic process | + | 4.00e-04 | 4.97e-02 |
|  |  | GO:0006555 | Methionine metabolic process | + | 2.63e-04 | 3.43e-02 |
|  |  | GO:0046942 | Carboxylic acid transport | + | 2.14e-05 | 4.64e-03 |
|  |  | GO:0019693 | Ribose phosphate metabolic process | + | 2.79e-04 | 3.54e-02 |
|  |  | GO:0006753 | Nucleoside phosphate metabolic process | + | 7.95e-05 | 1.34e-02 |
|  |  | GO:0042254 | Ribosome biogenesis | + | 9.32e-05 | 1.43e-02 |
|  |  | GO:0006897 | Endocytosis | - | 6.73e-06 | 1.95e-03 |
|  |  | GO:0032197 | Transposition, RNA-mediated | - | 1.25e-04 | 1.72e-02 |
|  | Molecular Function | GO:0046943 | Carboxylic acid transmembrane transporter activity | + | 2.09e-05 | 2.66e-02 |
|  |  | GO:0008514 | Organic anion transmembrane transporter activity | + | 5.63e-05 | 2.66e-02 |
|  |  | GO:0043167 | Ion Binding | + | 9.12e-05 | 4.63e-02 |
|  |  | GO:0003824 | Catalytic activity | + | 3.07e-06 | 7.79e-03 |
|  | Cellular Component | GO:0110165 | Cellular anatomical entity | + | 1.03e-05 | 1.09e-02 |

The *sam2Δ/sam2Δ* mutant strain had 876 genes with different expression patterns from wildtype baseline expression. This differentially expressed gene list consisted of 405 upregulated genes and 471 downregulated genes at the same cutoff values (p-value ≤ 0.05, 0.67 ≤ |FC| ≥ 1.5). A list and description of all upregulated and downregulated genes can be found in Supplementary Table 1. Gene ontology analysis of all DEGs from *sam2Δ/sam2Δ* cells revealed several enriched terms across the three domains (Table 1). Broadly, the deletion of both copies of *sam2* leads to differential expression of genes involved in the biological process of amino acid and deoxynucleoside triphosphate (dNTP) metabolism, while enriched molecular functions included processes involved in transmembrane transport, ion binding, protein and xenobiotic metabolism, and catalytic activity. One protein AdoMet-dependent methyltransferase gene was found on the upregulated gene list: *RKM4* (FC = 1.75, p-value =0.00565), along with one AdoMet-dependent methyltransferase gene whose encoded protein has an unknown substrate: *CRG1* (FC = 2.48, p-value = 0.00005), and one AdoMet-dependent small molecule methyltransferase: *TMT1* (FC = 2.48, p-value = 0.0003). Fifteen AdoMet-dependent methyltransferase genes were found on the downregulated gene list in this strain. There were two lipid AdoMet-dependent methyltransferase genes [*ERG6* (FC = 0.58, p-value = 0.04195), *OPI3* (FC = 0.54, p-value = 0.01215)], six protein AdoMet-dependent methyltransferase genes [*TAE1* (FC = 0.45, p-value = 0.0003), *EFM3* (FC = 0.52, p-value = 0.005), *EFM6* (FC = 0.63, p-value = 0.03995), *DPH5* (FC = 0.62, p-value = 0.03295), *PPM2* (FC = 0.48, p-value = 0.01045), and *NOP1* (FC = 0.45, p-value = 0.0023)], three rRNA AdoMet-dependent methyltransferase genes [*BUD23* (FC = 0.36, p-value = 0.00055), *NOP2* (FC = 0.50, p-value = 0.0057), and *EMG1* (FC =0.53, p-value = 0.0075)], and four tRNA AdoMet-dependent methyltransferase genes [*TRM5* (FC = 0.50, p-value =0.00645), *TRM9* (FC = 0.58, p-value = 0.0088), *TRM13* (FC = 0.62, p-value = 0.0225), and *TRM112* (FC = 0.48, p-value = 0.00075)] (Fig 1B).

In comparing the differentially expressed genes from each homozygous knockout strain, 59 genes (6.2%) were present on both lists. 44 of these 59 genes showed the same expression pattern in each mutant strain (26 with decreased expression, 18 with increased expression) whereas 15 genes showed opposite expression patterns between the two mutant strains (11 *sam1Δ/sam1Δ* decreased but *sam2Δ/sam2Δ* increased, and 4 *sam1Δ/sam1Δ* increased but *sam2Δ/sam2Δ* decreased) (Fig 2). A list of these genes can be found in Supplementary Table 3. Our previous research has revealed opposing patterns of genome stability due to altered AdoMet abundance in *sam1Δ/sam1Δ* and *sam2Δ/sam2Δ* mutant strains [2]. If the underlying cause of this genome stability is due to gene expression changes in these mutant strains, we would not expect genes with the same directionality of change in their expression profiles to be causatively linked. Rather, the 15 genes showing inverse expression patterns between our mutants have more potential to be driving the opposite impacts on genome stability, if this phenotype is driven solely by gene expression differences. Or, if there are multiple factors beyond just gene expression involved, then we might consider genes that have altered expression in one, but not both mutants, to be players of interest in the genome stability phenotypes.

**Fig 2.**
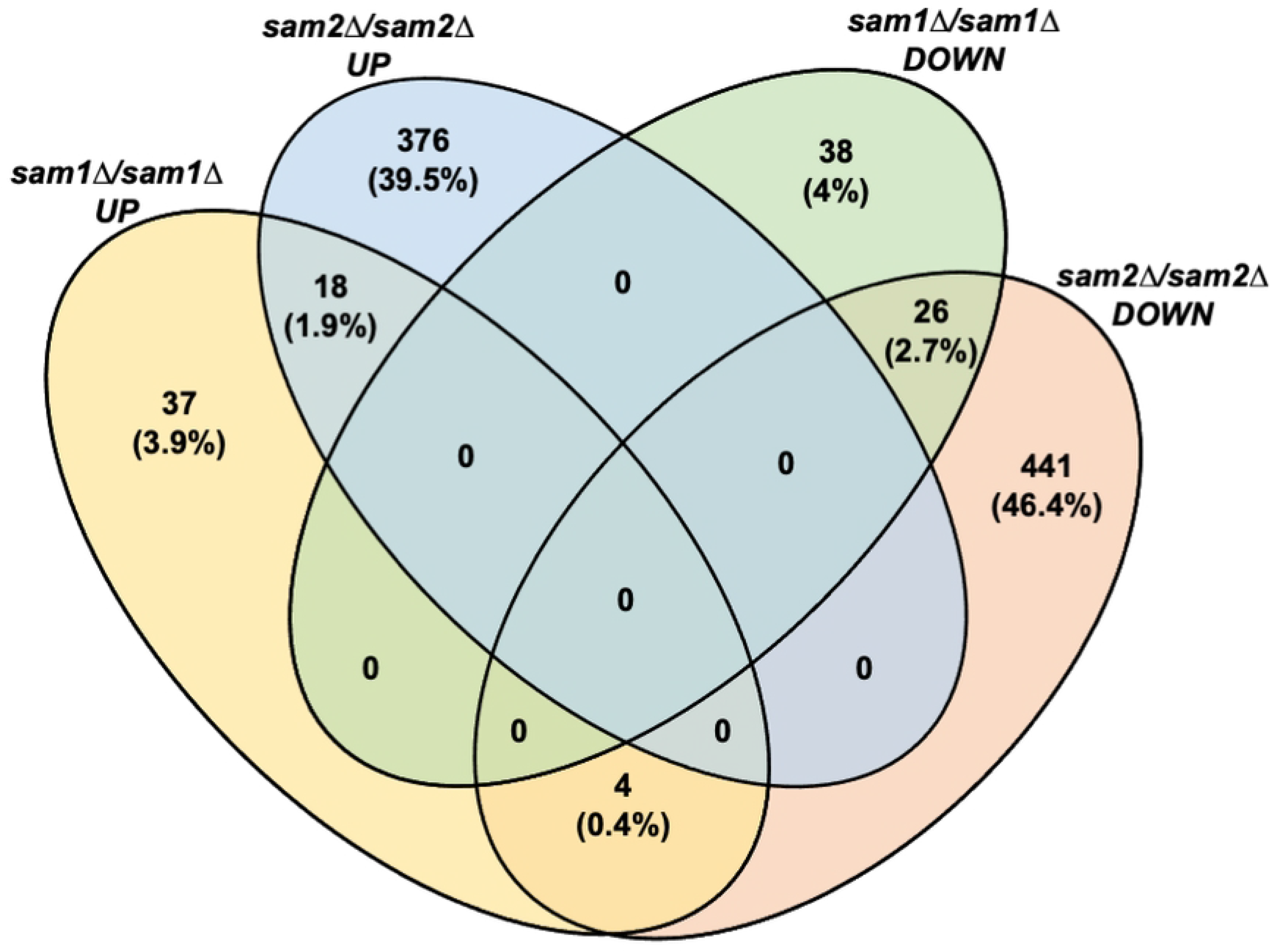
Comparison of DEGs found in *sam1Δ/sam1Δ and sam2Δ/sam2Δ* mutant strains. Venn diagram, created with Venny, showing the overlap of differentially expressed genes from the homozygous knockout mutant strains. Different categories show where the same genes were altered in their expression in the same direction in the mutant strains and where the *SAM* gene deletions had opposite impacts on the expression of the same genes.

### Phenotypic Microarray

We utilized 15 Phenotypic Microarray (PM) plates to characterize phenotypic growth differences in our mutant strains (PM1-10, PM21D, PM22D, PM23A, PM24C, and PM25D) [109]. There were 1440 total wells with 13 serving as either positive or negative controls. Several wells contained the same substrate but at varying concentrations. With those instances counting as a single condition, and controls removed, our wildtype and mutant strains were exposed to a total of 1005 completely different test conditions during growth. When analyzing growth to then define different phenotypes, three parts of the growth curves were assessed as was done in Fernandez-Ricaud et al. 2007 [112]. These parts include the adaptation time (time to start of growth), rate of growth, and efficiency of growth (time to saturation and maximum OD) (Fig 3A).

**Fig 3.**
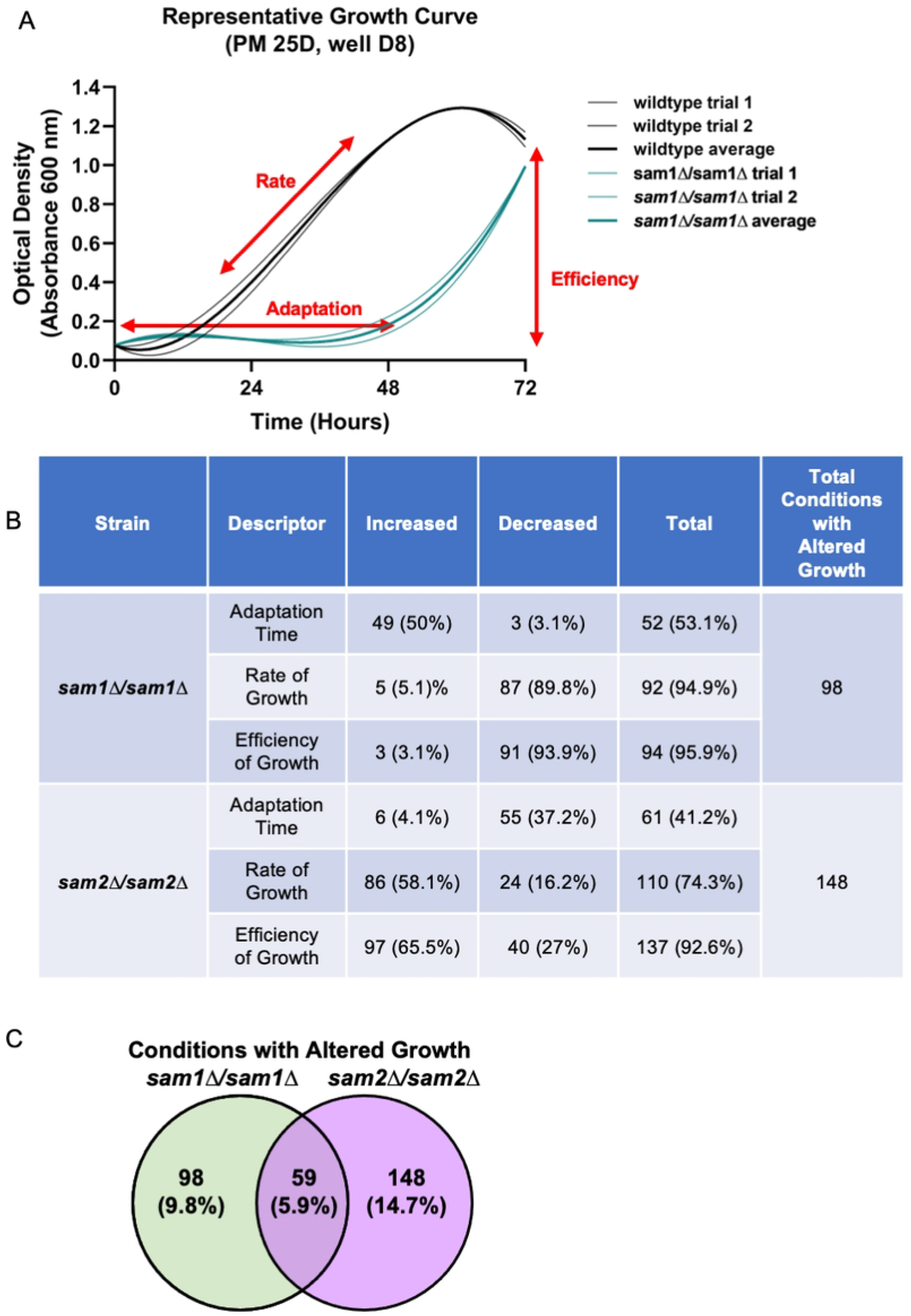
Characterization of Phenotypic Microarray results from our strains of interest. (A) representative growth curve showing growth patterns of biological replicates of wildtype (black) and *sam1Δ/sam1Δ* (green) strains. Red arrows indicate the application of growth metric descriptors as per Fernandez-Ricaud et al [112]. (B) Growth patterns were analyzed for each condition and growth descriptors were applied where mutant strains showed differences. Graphs were not interpreted if biological replicate data for each strain type were not consistent. A single condition could be counted for more than one growth descriptor category, as seen for *sam1Δ/sam1Δ* that exhibit an increased adaptation time and lower efficiency of growth in Hygromycin B. (C) Venn diagram showing the overlap of PM conditions that elicited altered growth patterns in the homozygous knockout mutant strains. While 59 conditions resulted in altered growth patterns for both the *sam1Δ/sam1Δ* and *sam2Δ/sam2Δ* mutant strains, the growth pattern changes were frequently not identical.

The *sam1Δ/sam1Δ* mutant strain, which exhibits increased AdoMet levels, showed patterns of growth that differed from the wildtype strain in 146 total wells that encompassed 98 different conditions tested, representing altered growth in 9.75% of conditions tested (Supplementary Table 4). For these conditions that had differing growth, the largest group (95.9%) impacted efficiency, whereas 94.9% exhibited altered rate of growth, and 53.1% impacted adaptation time (Fig 3B). The *sam2Δ/sam2Δ* mutant strain showed altered growth in 206 total wells that encompassed 148 different conditions (Supplementary Table 4). Thus, the full loss of *sam2* expression, which results in decreased AdoMet levels, has wider phenotypic impacts changing growth patterns in 14.7% of conditions tested. In the conditions that had differing growth, the largest group (92.6%) impacted growth efficiency, whereas 74.3% had altered rates of growth and only 41.2% differed in their adaptation time (Fig 3B).

Fifty-nine different conditions were identified as impacting growth in both homozygous knockout mutant strains (Fig 3C). Of these 59 conditions, 16 resulted in growth patterns of a similar nature in both the *sam1Δ/sam1Δ* and *sam2Δ/sam2Δ* mutant strains compared to wildtype; D-galactose, D-xylose, D-fructose, a-D-glucose, 5-keto-D-gluconic acid, 6% NaCl + ectoine, 6% NaCl + octopine, EDTA, bleomycin, potassium chromate, isoniazid, sodium fluoride, ibuprofen, blasticidin S, D-glucosamine, and L-lyxose. We therefore hypothesize that the pathways impacted by these conditions are not related to the increased and decreased AdoMet levels or increased and decreased chromosome stability seen in the *sam1Δ/sam1Δ* and *sam2Δ/sam2Δ* strains respectively, since the growth patterns are identical despite the different mutations. There were 21 different conditions that elicited growth patterns that were opposite in nature between the *sam1Δ/sam1Δ* and *sam2Δ/sam2Δ* mutant strains. The mutant strains showed opposite altered growth patterns to varying concentrations of NaCl, 4% sodium formate, pH 4.5 + L-ornithine, pH 4.5 + D,L-a-amino-n-butyric acid, protamine sulfate, magnesium chloride, sodium selenite, L-arginine hydroxamate, 3-amino-1,2,4-triazole, doxycycline, cobalt(II) chloride, sodium cyanide, sodium thiosulfate, sodium metasilicate, 6-azauracil, cisplatin, aluminum sulfate, fluconazole, miconazole, L-glutamic acid g-hydroxamate, and chloroquine. In these 21 conditions, the full loss of *sam1*, with its resulting increase in AdoMet levels, resulted in slower growth rates and lowered efficiencies, whereas the full loss of *sam2*, with its resulting decreased AdoMet levels, resulted in greater growth rates which sometimes were also accompanied with better growth efficiencies. These conditions that impact both strains, but in non-identical ways, as well as those conditions that impact one, but not the other mutant strain, are of interest, as the pathways that are impacted might underlie the previously observed chromosome stability changes which also have opposite patterns.

Looking for overarching patterns across all wells that resulted in altered growth, we characterized what percentage of 1005 different conditions tested fit into distinct mechanism of action categories. We then determined which of the 98 conditions where we saw altered growth in the *sam1Δ/sam1Δ* mutant cells and the 148 conditions where we saw altered growth in the *sam2Δ/sam2Δ* mutant cells, fit in these categories and calculated percentages to determine any areas of overrepresentation. We saw that 21.4% of the conditions where *sam1Δ/sam1Δ* mutant cells showed altered growth had a mechanism of action relating to toxic ions (2.9% of conditions in the PM wells tested), 8.1% related to protein synthesis (1% of PM wells tested), 7.1% related directly to membrane stability (1.6% of PM wells tested), 17.3% related to osmolarity/osmotic sensitivity (7.6% of PM wells tested), and 3% related to tRNA synthetases (0.5% of PM wells tested). The conditions that elicited altered growth in the *sam2Δ/sam2Δ* mutant cells could be broadly categorized as having mechanisms of action of which 4.1% related to protein synthesis (1% of PM wells tested), 6.8% related directly to membrane stability (1.6% of PM wells tested), 11.5% related to toxic ions (2.9% of PM wells), 4.6% related to respiration (0.7% of PM wells), 26.4% related to osmolarity/ osmotic sensitivity (7.6% of PM wells tested), and 2.7% related to tRNA synthetases (0.5% of PM wells tested).

### Linking Phenotypic Changes with Underlying Gene Expression Differences

The use of RNA-seq and the PM allowed broad characterization of differences in our *sam1Δ/sam1Δ* and *sam2Δ/sam2Δ* mutant strains. While these experiments uncovered more information about cellular changes due to deletions of copies of the S-Adenosylmethionine synthetase genes, *SAM1* and *SAM2*, additional work was needed to try to identify how these mutations, and the resultant alterations in AdoMet concentration, led to the various differences that were observed. Therefore, we next sought to link the data collected from our RNA-Seq experiments to the phenotypic differences from the PM data, to define pathway alterations that led to significant growth changes for our mutants. To do this we looked at the mechanisms of action of the conditions of a particular PM well and then sought to map the affected pathways. Next, we overlayed our Differentially Expressed Gene (DEG) data to see if the *sam1Δ/sam1Δ* or *sam2Δ/sam2Δ* deletions altered expression of any genes in these pathways. Finally, we sought to determine if the changes in expression of impacted pathway genes, impacts to the methyl cycle, altered AdoMet-dependent methyltransferase expression, or AdoMet concentration differences/availability, could explain the observed growth differences of the strains.

#### A. Lack of *sam2* confers resistance to anti-fungal azole drugs, while *sam1*-deficient cells show increased sensitivity

The 15 PM plates we used contain 3 different azole fungicides (Supplementary Table 5). These conditions and their BiOLOG reported mechanisms of action are fluconazole - “inhibits Lanosterol 14 α-demethylase” (PM 24C G09-12), miconazole - “triazole fungicide” (PM24C H09-12), propiconazole - “triazole fungicide” (PM24C H01-04). We observed growth differences in our *sam1Δ/sam1Δ* and/or *sam2Δ/sam2Δ* mutant strains in all three of these conditions. Fig 4 shows the growth curves of the wildtype, *sam1Δ/sam1Δ* and *sam2Δ/sam2Δ* strains in response to these different azoles from the PM plates. Where multiple concentrations were tested in different PM wells, data from a representative well is shown. (For all growth curves see Supplementary Tables 6 and 7).

**Fig 4.**
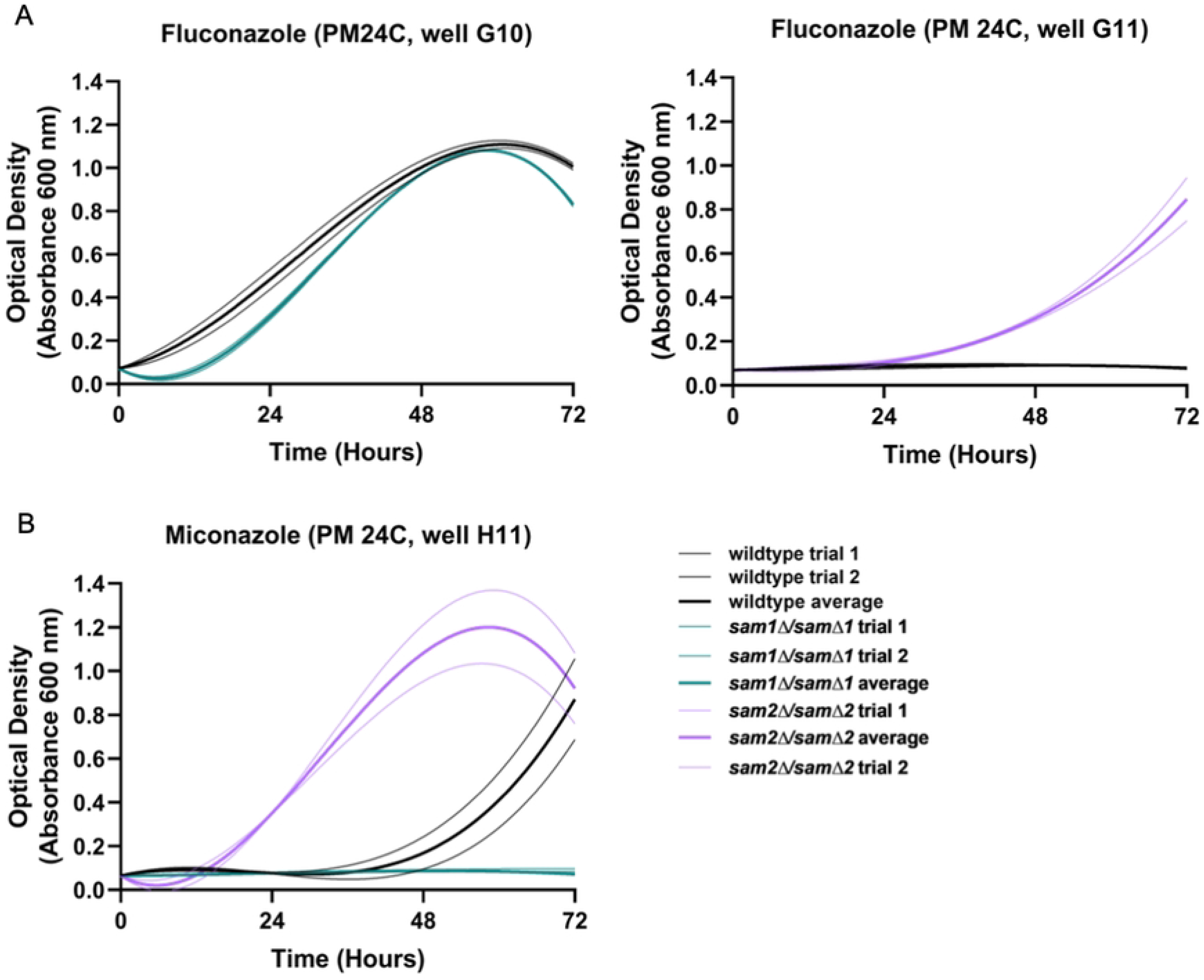
Growth curves of wildtype, *sam1Δ/sam1Δ* and *sam2Δ/sam2Δ* cells in the presence of (A) Fluconazole and (B) Miconazole. Growth curves generated from timepoint data of the Phenotypic Microarray experiments. The independent trials for each strain are shown in a lighter weight line, while the average is shown in the same color in bold. (A) Growth curves are shown for wildtype and *sam1Δ/sam1Δ* cells in the third highest concentration of Fluconazole (G10) and wildtype and *sam2Δ/sam2Δ* cells in the second highest concentration (G11). (B) Growth curves are shown for wildtype, *sam1Δ/sam1Δ, sam2Δ/sam2Δ* cells in the second highest concentration of Miconazole (H11).

In PM plate 24C well G10, containing fluconazole, the *sam1τι/sam1τι* mutant cells exhibited decreased growth rate and increased adaptation time, compared to the wildtype, with growth beginning around 12 hours. This indicates that these cells are sensitive at this concentration (Fig 4A). In PM plate 24C well G11, the *sam211/sam211* mutant cells started to grow at the 24-hour timepoint and maintained increased growth the remainder of the experiment (Fig 4A). Compared to wildtype cells, *sam211/sam211* mutant cells exhibited an increased rate and efficiency of growth indicating increased resistance to this condition starting at 24 hours (Fig 4A). In PM plate 24C well H11 containing Miconazole, wildtype cells began to grow around the 36-hour timepoint and grew fastest between 48 and 72 hours (Fig 4B). The *sam111/sam111* mutant cells were unable to grow, indicating sensitivity at this concentration (Fig 4B). The *sam211/sam211* mutant cells started to grow at the 12-hour timepoint and grew fastest between 12 and 48 hours (Fig 4B). Compared to wildtype cells, *sam211/sam211* mutant cells exhibited a decreased adaptation time and increaseFig14d growth efficiency (Fig 4B). In PM plate 24C wells H1-H4 containing Propiconazole, the growth patterns do show differences between the wildtype and mutant strains, however we do not see enough consistency in growth trends between biological replicates of each strain to support further discussion of this condition here.

Between the cell wall and the cytosol of *S. cerevisiae* lies the plasma membrane, which is comprised of a lipid bilayer with a wide array of lipids and proteins that work together to maintain membrane integrity and monitor membrane permeability [113]. Sterol products serve as important structural components of the plasma membrane [114]. Cholesterol is the main sterol present in eukaryotic plasma membranes, except fungal plasma membranes, whose crucial component is ergosterol as the main sterol, along with small amounts of zymosterol and other sterols [115,116]. Much like cholesterol in mammalian cells, ergosterol functions to maintain membrane integrity by contributing to proper structure, permeability, and fluidity [116,117]. In *S. cerevisiae*, the ergosterol biosynthesis pathway has three sections named the mevalonate pathway, the late pathway, and the alternate pathway (Fig 5). Many anti-fungal agents target the ergosterol biosynthesis pathway, including the azoles [118].

**Fig 5.**
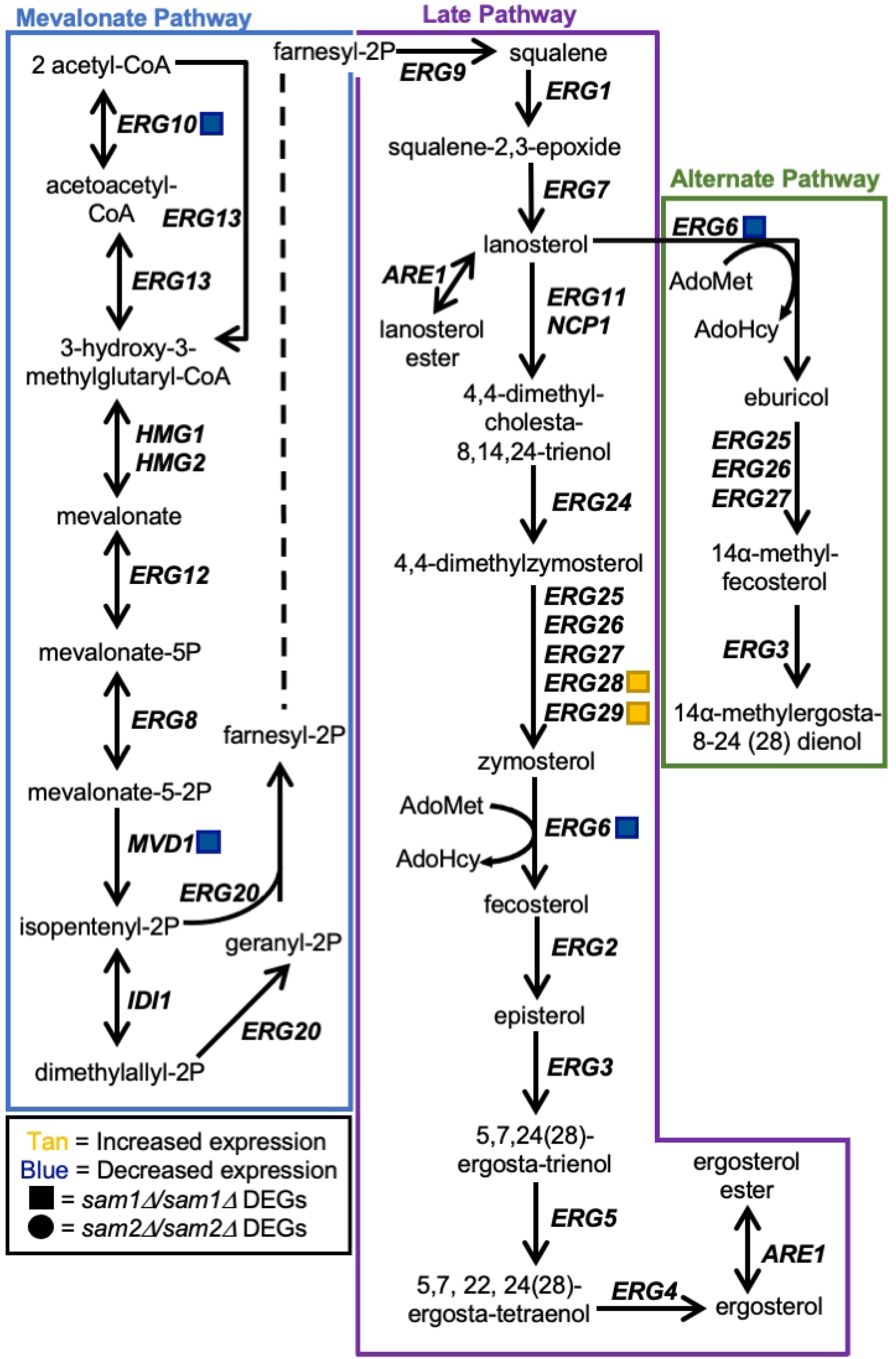
Ergosterol Biosynthesis Pathway. Production of ergosterol in *S. cerevisiae* begins with two acetyl-CoA molecules and produces additional sterols through the interconnected mevalonate, late, and alternate pathways. The output of the mevalonate pathway, farnesyl-2P, is the starting substrate for the late pathway. The gene names that encode the enzymes that can function in compound conversion in the pathway are given by their standard names. *sam2Δ/sam2Δ* DEGs are represented by red (increased expression) and green (decreased expression) squares. RNA-sequencing did not reveal any differentially expressed genes within this pathway between *sam1Δ/sam1Δ* and wildtype.

Azoles are synthetic organic compounds that consist of five-membered rings containing one nitrogen atom along with another non-carbon atom [119]. The development of azoles were prompted as many species of pathogenic fungi cause severe plant and human disease [120,121]. There are two subclasses of azoles: imidazoles and triazoles [122,123]. Fluconazole, propiconazole, and miconazole, are triazoles that selectively target lanosterol 14α-demethylase, which is a rate-limiting enzyme encoded by the gene *ERG11*. Lanosterol 14α-demethylase is a cytochrome P450-dependent enzyme responsible for the conversion of lanosterol to 4,4-dimethylcholesta-8,14,24-trienol within the late pathway [124–127]. These azoles selectively inhibit lanosterol 14α-demethylase through binding: for instance, the azole nitrogen of fluconazole binds to the heme group of the enzyme, thus preventing the enzyme’s ability to bind to the proper substrate and produce products that are used in the late pathway [128,129]. As a result, cells use the alternate pathway to produce 14α-methylergosta-8,24(28)-dienol which compensates for lack of ergosterol, but is also toxic to the cell in high enough concentrations [129–134].

Although 14α-methylergosta-8,24(28)-dienol is a toxic 14α-methyl sterol, other 14α-methyl sterols are compatible with growth, such as lanosterol and 14α-methylfecosterol [135– 137]. The toxic 14α-methyl sterols are problematic because accumulation of these aberrant sterols interrupts proper packing of acyl chains in the phospholipid bilayer and they abnormally interact with other components of the membrane, causing a disruption of ideal structure and functionality [138,139]. While miconazole has actions similar to that of the other triazoles, miconazole is more complex as multiple mechanisms of action exist. Beyond inhibiting lanosterol 14α-demethylase, miconazole treatment results in increased intracellular reactive oxygen species (ROS) due to altered peroxide and catalase enzyme activity, interference with triglyceride synthesis, inhibition of the utilization of glucose, and membrane damage [124,140–143]. Additional mechanisms aside, by inhibiting lanosterol 14α-demethylase, triazoles prevent the production of ergosterol and cell membrane integrity becomes compromised, due to the accumulation of the toxic dienol which inhibits fungal growth [117,132,135–137]. As seen in Fig 4, strains homozygously deleted for *sam2* are resistant to these azole drugs as compared to wildtype. Both the late and alternate pathways of the ergosterol biosynthesis pathway involve an AdoMet-dependent methyltransferase, called sterol C-24-methyltransferase, encoded by the *ERG6* gene. Erg6 in the late pathway is responsible for the conversion of zymosterol into fecosterol through methylation. Fecosterol is then processed through the late pathway to be converted to ergosterol as the final product. Erg6 in the alternate pathway is responsible for the methylation of lanosterol into eburicol. Eburicol is ultimately converted to the toxic sterol, 14α-methylergosta-8,24(28)-dienol through a series of two reactions; one involving the enzyme complex (Erg25, Erg26, Erg27) that removes 2 methyl groups, while the other involves Erg3 which creates a double bond [133].

Through investigation of the full list of DEGs from the *sam2Δ/sam2Δ* strain, we were able to identify *ERG10*, *MVD1*, *ERG6*, *ERG28*, and *ERG29* of the ergosterol synthesis pathway as showing altered expression. The expression genes of the DEGs in the *sam2*11*/sam2*11 mutant strain are: *ERG10* (fold change of 0.55, p-value=0.04245), *MVD1* (fold change of 0.55, p-value=0.03155), *ERG6* (fold change of 0.58, p-value=0.04195), *ERG28* (fold change of 1.56, p-value = 0.0477), and *ERG29* (fold change of 1.59, p-value = 0.03975) (Fig 5).

The decrease in *ERG6* expression in the *sam2Δ/sam2Δ* strain appears to lessen the input into the alternate pathway, while the upregulation of *ERG28* and *ERG29* in the late pathway would appear to increase zymosterol production. This increase in expression of *ERG28* and *ERG29* could benefit the cell two ways; first, being downstream of two genes with reduced expression in the mevalonate portion of the pathway, their increased expression could serve to more rapidly allow these cells to achieve pathway functionality at or near wildtype, or higher, levels for zymosterol production and second, *ERG6* functions directly downstream of these genes and increasing the substrate when the enzyme is limited may increase conversion to fecosterol overall to facilitate completion of the late pathway to result in ergosterol production at more normal levels. In the *sam2Δ/sam2Δ* strain, we have previously shown that these cells have a significant reduction in their AdoMet levels, which could be hypothesized to further limit methylation by Erg6 through the reduced methyl donor availability. Thus, this decreased expression of *ERG6*, and the reduced concentration of AdoMet, can explain the resistance to fluconazole, propiconazole, and miconazole. While Erg6 functions in both the late and alternate pathways it’s reduced activity in the alternate pathway results in less 14α-methylergosta-8,24(28)-dienol production, which then cannot build up to a toxic level and these cells are resistant to these drugs as compared to the wildtype cells.

Unlike in the *sam2Δ/sam2Δ* strains, we saw no changes in expression in any of these genes in this pathway due to the full loss of *sam1*. Further, we have characterized *sam1*-deficient strains as having an increased concentration of AdoMet [2]. This abundance of methyl-donor could be hypothesized to explain the increased sensitivity of the *sam1Δ/sam1Δ* cells to these azole drugs in that while expression of the *ERG6* gene is not increased, it’s rate of activity may be enhanced by the increased AdoMet. As seen for fluconazole, *sam111/sam111* cells show consistent growth patterns with wildtype strains, whereas in miconazole they take longer to adapt and have reduced efficiency. Therefore, when Erg11 is targeted by fluconazole or miconazole, cells shift to the alternate pathway in an attempt to survive, causing their sensitivity in the same manner as is seen in wildtype strains. With the miconazole concentrations tested we may have also captured increased sensitivity of the *sam1*-deficient cells that may indicate that they produce the toxic dienol faster than wildtype cells due to the increased abundance of the AdoMet methyl donor for the methylation reaction required for this transition; or the increased sensitivity of the *sam1Δ/sam1Δ* cells to miconazole versus the other azoles may reflect on miconazole’s additional mechanisms of action on cells.

#### B. Lack of *sam2* confers resistance to cisplatin

The PM plates contain cisplatin (PM24C G1-4), a known chemotherapeutic. The reported mechanism of action for cisplatin was classified by BiOLOG as “DNA damage, crosslinker”. We observed phenotypic growth differences in both of our mutant strains in this condition. Fig 6 shows the growth curves of the wildtype, *sam111/sam111*, and *sam2Δ/sam2Δ* strains in response to cisplatin in the PM plates. Multiple concentrations were tested in different PM wells and data from a representative well is shown for each mutant. (For all growth curves see Supplementary Tables 6 and 7).

**Fig 6.**
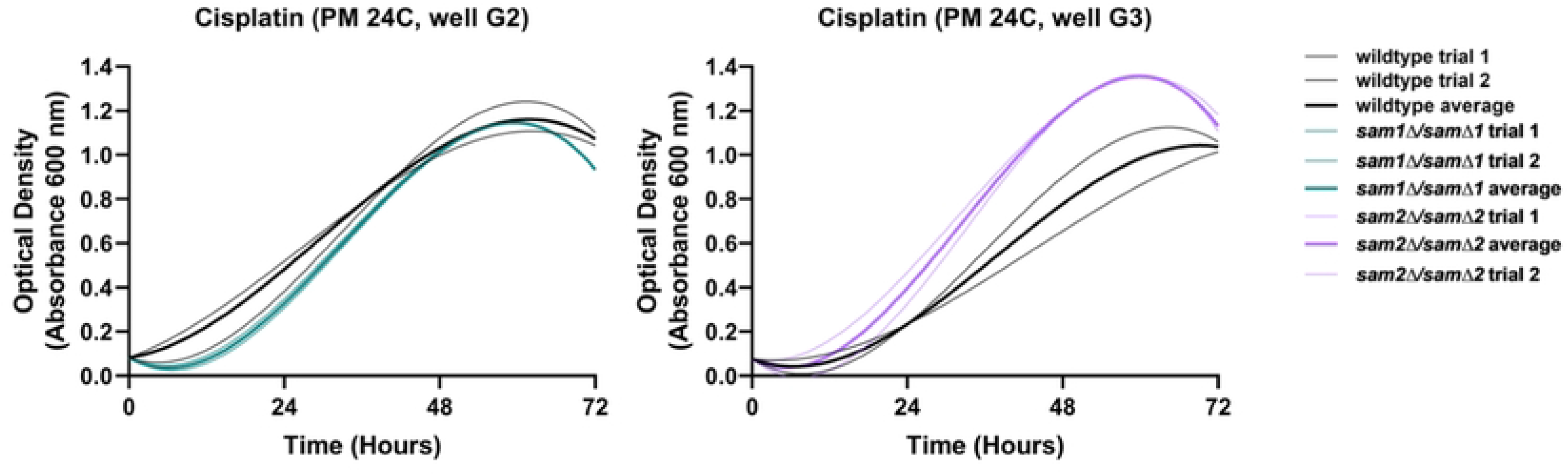
Growth curves of wildtype, *sam1Δ/sam1Δ* and *sam2Δ/sam2Δ* cells in the presence of Cisplatin. Growth curves generated from timepoint data of the Phenotypic Microarray experiments. The independent trials for each strain are shown in a lighter weight line, while the average is shown in the same color in bold. Growth curves are shown for wildtype and *sam1Δ/sam1Δ* cells in the third highest concentration of Cisplatin (G2) and wildtype and *sam2Δ/sam2Δ* cells in the second highest concentration (G3).

In PM plate 24C well G2, the *sam111/sam111* mutant cells have an increased adaptation time compared to wildtype, with growth beginning around 12 hours, and exhibited lower growth rate (Fig 6). In PM 24C well G3, the *sam211/sam211* mutant cells exhibited decreased adaptation time, with growth beginning around 12 hours, and increased growth rate and efficiency in comparison to wildtype, on average (Fig 6).

Cisplatin is a strong platinum-based drug, with genotoxic, cytostatic, and cytotoxic abilities [144]. Cisplatin is taken up by *S. cerevisiae* through utilization of the copper transporter genes *CTR1*, *CTR2, CTR3,* and *FET4* (Fig 7), each of which have different affinities for copper, but can bind to platinum-based drugs as well due to the presence of a metal binding motif [145]. Once in the cell, copper is utilized in the mitochondria for enzyme complexes and ATP production, in the cell wall for cellular integrity, or sent to organelles for functions such as cell signaling and cellular proliferation [146]. Cisplatin, on the other hand, causes damage to DNA by forming covalent bonds between the platinum ion and the N7 position of purine bases, which in turn forms a 1,2- or 1,3-intrastrand crosslink [147]. Consequently, the DNA becomes “clunky” and is unable to be replicated or transcribed, which may cause cell cycle arrest and apoptosis [147,148]. Previous studies have found that the effects of cisplatin can be ultimately reversed through multiple DNA repair mechanisms, including Nucleotide Excision Repair (NER), MisMatch Repair (MMR), and Recombinational Repair (RR) [149]. NER is a mechanism in which the cell is able to remove DNA lesions caused by mutagens, through the unwinding and excision of the damaged DNA, followed by gap filling of missing base pairs by DNA polymerase and ligation by DNA ligase [150]. MMR is similar to NER, but it corrects only small mismatches in DNA instead of entire lesions. MMR takes place through recognition of mismatched base pairs in DNA caused by mutagens or replicative mistakes, followed by excision of a small segment of DNA, and finally the correct bases are added to the gap by DNA polymerase [151]. RR occurs in eukaryotes only, and happens when a section of damaged DNA is replaced with an exact copy of non-damaged DNA by a strand exchange protein (most commonly of the RAD family of proteins), followed by ligation by DNA ligase [152]. Cells also contain several membrane-spanning proteins that have the capability to pump out toxic metals [153]. We therefore envisioned that cell sensitivity or resistance to cisplatin could be impacted through changes in the uptake of the drug, reaction/repair to damage caused by the drug, or altered export of the drug.

**Fig 7.**
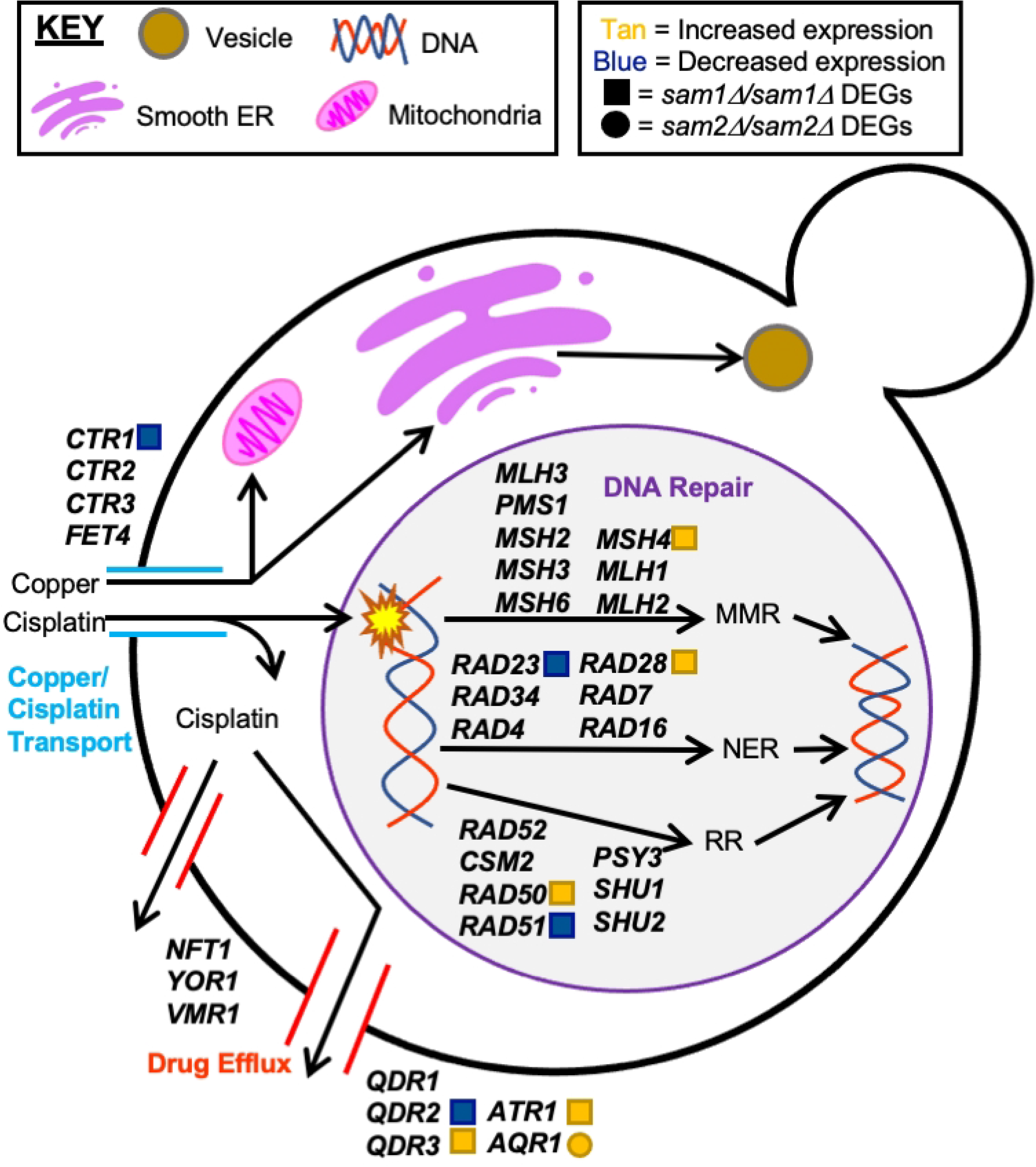
Copper/Cisplatin Transport Pathway, DNA Repair Pathways, and Drug Efflux Pathways. Uptake of copper and cisplatin in *S. cerevisiae* begins with import by membrane-associated transport proteins. Copper aids in several cellular processes in specialized organelles, such as the mitochondria, smooth endoplasmic reticulum, and vesicles. Cisplatin causes DNA damage, that can be subsequently repaired by DNA repair pathways: MMR, NER, or RR. Cisplatin can be exported out of the cell by specialized drug efflux pumps. The gene names that encode the enzymes that can function in compound conversion in the pathway are given by their standard names. *sam2Δ/sam2Δ* DEGs are represented by red (increased expression) and green (decreased expression) squares. *sam1Δ/sam1Δ* DEGs are represented by red (increased expression) and green (decreased expression) circles.

Through investigation of the full list of DEGs from the *sam1Δ/samΔ1* and *sam2*11*/sam2*11 mutant strains, we were able to identify *CTR1* of the copper/cisplatin transport pathway, *MSH4, RAD50, RAD51, RAD28,* and *RAD23* of the DNA repair pathway, and *QDR2, QDR3, AQR1,* and *ATR1* of the drug efflux pathway (Fig 7) as differentially expressed. We explored differentially expressed genes in our mutant strains within these pathways to determine if the observed phenotypic growth differences could be explained at this level of regulation.

Within the copper/cisplatin transport pathway (Fig 7), the *sam2Δ/sam2Δ* mutant cells exhibited decreased expression of *CTR1* (fold change of 0.52, p-value=0.00285). As previously mentioned, Ctr1 is responsible for import of copper, and is the highest-affinity transporter of all the copper transporters, but it can also bind to and import metal cation drugs, such as cisplatin. Previous studies have found that loss of *CTR1* results in almost complete resistance to cisplatin due to the lowered ability of cisplatin to be transported into the cell [145]. We hypothesize that because *sam2*11*/sam2*11 mutant cells have lowered Ctr1, that they can at least partially resist the deleterious effects of cisplatin due to decreased import and thus exposure of their DNA to the drug.

Within the DNA repair pathway, which includes NER, MMR, and RR, we observed several DEGs in the *sam2*11*/sam2*11 mutant strain (Fig 7). Within the NER pathway, we identified *RAD28* (fold change of 1.53, p-value=0.0308) and *RAD23* (fold change of 0.65, p-value=0.04125) (Fig 7). The MMR pathway contains one upregulated DEG, *MSH4* (fold change of 1.69, p-value=0.01675). Within the RR pathway, we found upregulation of *RAD50* (fold change of 1.69, p-value=0.01035), and downregulation of *RAD51* (fold change of 0.56, p-value=0.0092). The *sam2Δ/sam2Δ* mutant cells have both upregulated and downregulated genes within these pathways, so it is difficult to hypothesize an overall positive or negative impact, but rather we hypothesize that the growth changes we see in these cells is more likely due to the transport and efflux pathways of cisplatin.

Within the drug efflux pathway, three genes have been identified as responsible for exporting cisplatin only: *NFT1, YOR1,* and *VMR1,* each of which are part of the multidrug resistance-associated family of proteins (Fig 7) [154]. Although none of these genes are DEGs in our mutants, the *sam2Δ/sam2Δ* mutant cells do exhibit differentially expressed genes in a similar family of transporters, that export several drugs including cisplatin: we found downregulation of *QDR2* (fold change of 0.48, p-value=0.0007), and upregulation of *QDR3* (fold change of 1.76, p-value=0.00705) and *ATR1* (fold change of 1.66, p-value=0.01755) (Fig 7). The Qdr1, Qdr2 and Qdr3 proteins are multidrug/hydrogen ion antiporters, and are known to be responsible for cisplatin resistance [155]. Interestingly, *QDR1* has a paralog that arose via gene duplication, called *AQR1,* which encodes a protein able to pump out cisplatin, and confer resistance to this drug [156]. The *sam1Δ/samΔ1* mutant cells contain upregulated *AQR1* (fold change of 2.21, p-value=0.00005). *ATR1* also encodes a multidrug transporter, and upregulation of this gene has been found to correlate with cisplatin resistance as well [157]. We hypothesize that because the *sam2Δ/sam2Δ* mutant cells exhibit upregulation of two out of the four genes responsible for this family of multidrug export, that they are likely able to pump out more cisplatin than wildtype or *sam1Δ/samΔ1* mutant cells, therefore contributing to their overall increased growth observed in the PM plates for this condition.

We found no DEGS, aside from *AQR1,* present in the *sam1Δ/samΔ1* mutant strains in any of these pathways. The increased expression of *AQR1* could help explain why the *sam1*-deficient cells are able to reach the same growth saturation by 48-hours as the wildtype cells. However, the mechanisms responsible for the initial cisplatin sensitivity and longer adaptation time are not likely tied to altered import or DNA repair, and further research is warranted.

#### C. Lack of *sam2* leads to altered growth in conditions impacting glutathione and ROS

The PM plates contain nine different conditions that could impact Reactive Oxygen Species (ROS) levels and/or the glutathione (GSH) biosynthesis pathway, used to respond to ROS (Supplementary Table 5). These conditions can be grouped into two categories: drugs that increase ROS levels or decrease the cells’ ability to respond to ROS, and drugs that decrease ROS levels or increase the cells’ ability to respond to ROS. The conditions that increase ROS or decrease ROS response include sodium selenite (PM21D G01–04), methyl viologen (PM24C F5-8), urea hydrogen peroxide (PM22D D5-8), and thiourea (PM21D H5-8). The conditions that decrease ROS or increase ROS response include diamide (PM21D H1-4), and sodium thiosulfate (PM23A H9-12). The reported mechanisms of action were classified by BiOLOG as follows: sodium selenite-“toxic anion”, methyl viologen-“oxidizing agent”, urea hydrogen peroxide-“oxidizing agent”, thiourea-“membrane, chaotropic agent”, diamide-“oxidizes sulfhydryls, depletes glutathione”, and sodium thiosulfate-“toxic anion, reducing agent”. We observed growth differences in our *SAM* gene deleted strains in five of these conditions: sodium selenite, methyl viologen, thiourea, diamide, and sodium thiosulfate. [In the presence of diamide we observed growth differences, however, in researching its mechanism we found it impacts many cellular processes (oxidation, protein S-glutathionylation, translation, endoplasmic reticulum function, and cell wall integrity) therefore we cannot conclude that the growth changes we see are due to effects on GSH and/or ROS, and will not discuss this condition further [158–161].] Fig 8 shows the growth curves of the wildtype, *sam111/sam111*, and *sam2Δ/sam2Δ* strains in response to these compounds in the PM plates. Multiple concentrations were tested in different PM wells and data from a representative well is shown. (For all growth curves see Supplementary Tables 6 and 7).

**Fig 8.**
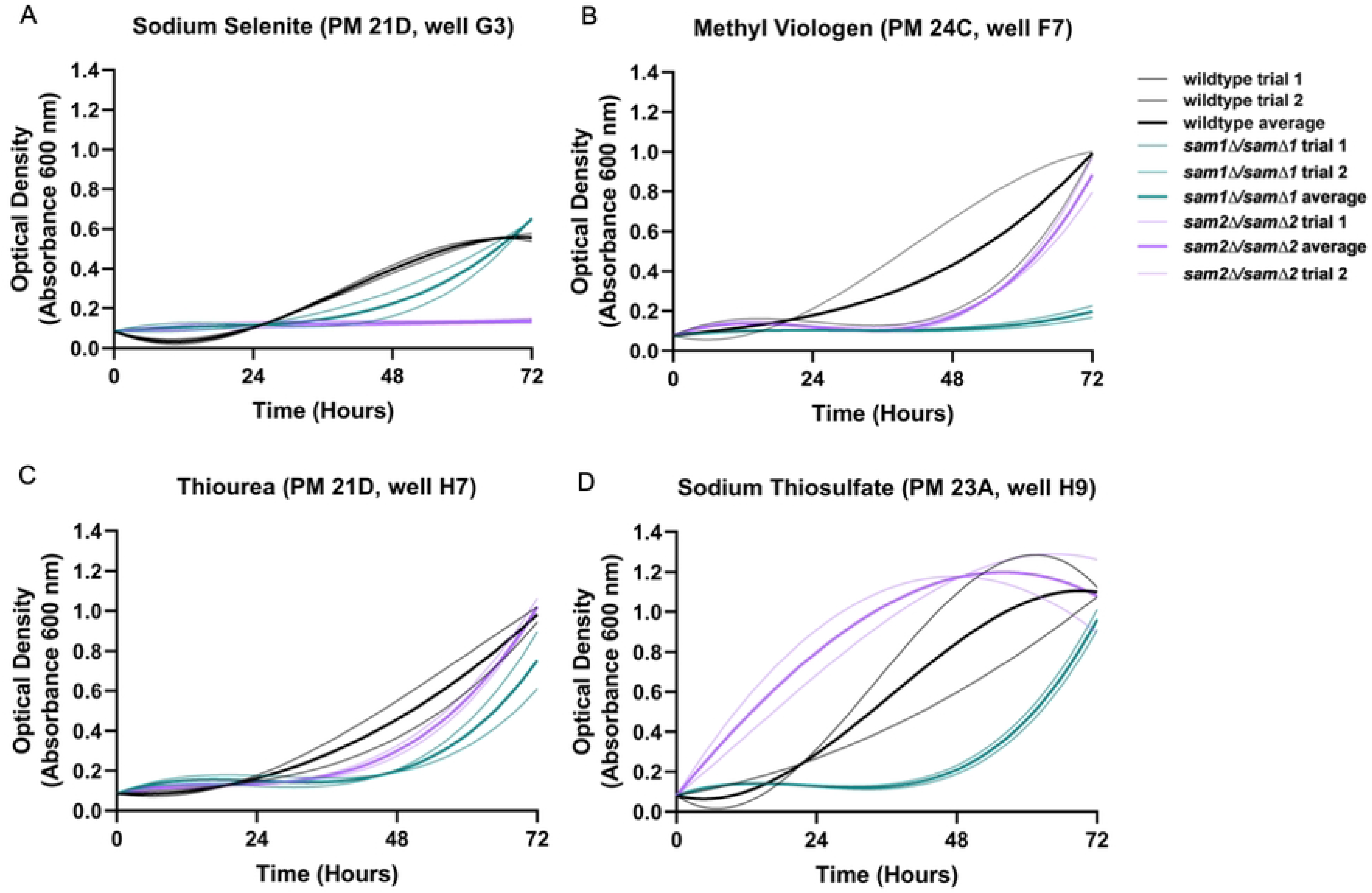
Growth curves of wildtype, *sam1Δ/sam1Δ* and *sam2Δ/sam2Δ* cells in the presence of (A) Sodium Selenite, (B) Methyl Viologen, (C) Thiourea, and (D) Sodium Thiosulfate. Growth curves generated from timepoint data of the Phenotypic Microarray experiments. The independent trials for each strain are shown in a lighter weight line, while the average is shown in the same color in bold. (A) Growth curves are shown for wildtype, *sam1Δ/sam1Δ, sam2Δ/sam2Δ* cells in the second highest concentration of Sodium Selenite (G3). (B) Growth curves are shown for wildtype, *sam1Δ/sam1Δ, sam2Δ/sam2Δ* cells in the second highest concentration of Methyl Viologen (F7). (C) Growth curves are shown for wildtype, *sam1Δ/sam1Δ, sam2Δ/sam2Δ* cells in the second highest concentration of Thiourea (H7). (D) Growth curves are shown for wildtype, *sam1Δ/sam1Δ, sam2Δ/sam2Δ* cells in the second highest concentration of Sodium Thiosulfate(H9).

In PM plate 21D well G3 containing sodium selenite, the *sam111/sam111* mutant cells show an increased adaptation time compared to wildtype but similar growth rate and efficiency (Fig 8A), whereas the *sam211/sam211* mutant cells exhibited no growth during the experiment (Fig 8A). In PM plate 24C well F7 containing methyl viologen and PM plate 21D well H7 containing thiourea, the *sam111/sam111* mutant cells exhibited decreased growth rate and efficiency and an increased adaptation time, whereas the *sam211/sam211* mutant cells exhibited no significant growth differences, compared to wildtype (Fig 8B & 8C). (The growth curve for *sam211/sam211* cells in methyl viologen is not considered significantly different from wildtype due to an overlap of growth patterns between one wildtype trial over the entire time course) In general, in the increased ROS/decreased ROS response conditions, the *sam111/sam111* mutant cells exhibit increased adaptation time and equivalent or slightly lowered growth rates and efficiencies compared to wildtype, whereas the *sam211/sam211* mutant cells exhibit either no growth at all (full sensitivity) or similar growth to wildtype.

In the decreased ROS/increased ROS response condition of sodium thiosulfate, in PM plate 23A well H9, the *sam111/sam111* mutant cells exhibit increased adaptation time and decreased growth efficiency (Fig 8D). At the same concentration, the *sam211/sam211* mutant cells exhibited increased growth rate and decreased adaptation time with growth beginning from the start (Fig 8D).

Sodium selenite and methyl viologen can both increase ROS levels and decrease the cells’ ability to combat ROS, while thiourea’s actions are concentrated on decreasing the cells’ ability to neutralize different ROS. Previous studies have determined that low concentrations of selenite are beneficial to *S. cerevisiae* by preventing spontaneous mutagenesis, but that high concentrations of selenite are toxic to cells [162,163]. Selenite reacts spontaneously with glutathione (GSH) in a five-step reaction mechanism [164–168]. The toxicity of selenite at high concentrations is seemingly caused by the transitory reaction intermediate hydrogen selenide (H_2_Se) [168]. It has been proposed that hydrogen selenide (H_2_Se) both oxidizes GSH and reacts with molecular oxygen in the cell to form ROS. Both reactions lead to oxidative stress due to an imbalance of GSH:GSSG. Previous research has found that with increasing concentrations of sodium selenite (Na_2_SeO_3_), both cell growth and GSH content decreased whereas selenide content, GSSG content, GSH peroxidase activity, and glutathione S-transferase (GST) activity increased [168,169]. Methyl viologen (MV), on the other hand, acts as an electron donor to GSSG, activating glutathione reductase to add two hydrogen groups onto GSSG and break the disulfide bond [170]. In plant cells, MV has been found to produce superoxide radicals that disrupt electron transfer [171]. These superoxide anions are a common precursor to ROS, and also have the ability to modify/deactivate cellular proteins, inhibit electron transport, and cause premature cell death [172,173]. Thiourea, another drug that induces ROS, does so by reducing GSH and NADPH until they are ultimately depleted in the cell [174]. Thiourea reacts with NADPH and oxygen to produce formamidine sulfenic acid, which catalyzes the oxidation of GSH to GSSG [175]. Without sufficient GSH and NADPH, the cell cannot neutralize ROS that are produced naturally elsewhere in the cell.

Sodium thiosulfate decreases ROS by readily donating electrons to the radical ROS species, inactivating them and rendering them harmless [176]. Previous research has found high concentrations of sodium thiosulfate to be toxic; however, as the wildtype and mutants grew, we conclude there are concentrations present in the PM plates that do not reach this level [177].

Through investigation of the full list of DEGs from the *sam2Δ/sam2Δ* and *sam1*11*/sam1*11 mutant strains, we were able to identify *MET6, SAM1, SAM2, SAH1, CYS3, CYS4, STR3, STR2, GSH1, GPX2,* and *DUG1 of* the glutathione biosynthesis pathway as showing altered expression. In the *sam111/sam111* mutant strain, the only gene in this pathway found to have differential expression was *SAM1* itself (whose absence is reflected as a significantly decreased DEG) (Fig 9).

**Fig 9.**
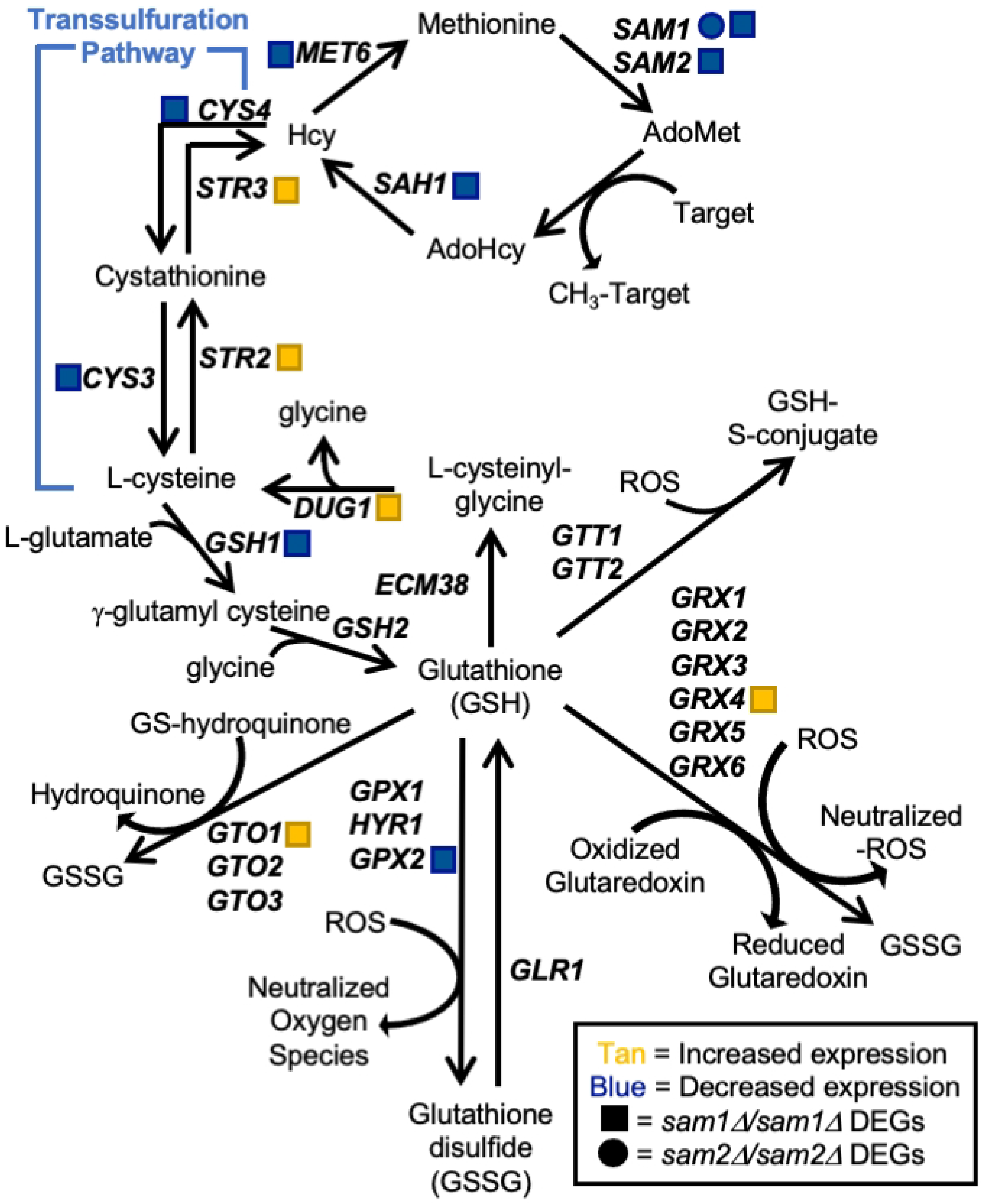
Glutathione Biosynthesis Pathway. Production of glutathione in *S. cerevisiae* requires the two component amino acids L-cysteine and glycine. L-cysteine is synthesized from L-homocysteine (Hcy), an intermediate of the methyl cycle. GSH1 and GSH2 are energy-requiring synthetases responsible for catalyzing the two consecutive steps of GSH biosynthesis. The peroxidases GPX1, HYR1, and GPX2 catalyze oxidation reactions whereas GLR1 is a reductase that catalyzes the reduction of GSSG to GSH to restore its antioxidative properties. Glutathione S-transferases (GSTs) are responsible for catalyzing the conjugation of toxic electrophiles to GSH in preparation for their removal from the cell. *sam2Δ/sam2Δ* DEGs are represented by red (increased expression) and green (decreased expression) squares. *sam1Δ/sam1Δ* DEGs are represented by red (increased expression) and green (decreased expression) circles.

GSH is an endogenously produced compound that possesses anti-oxidative properties that are important in protecting cells from oxidative stress. In its reduced form, GSH is a reactant needed for GSTs to neutralize ROS, such as hydroxyls and superoxides [178]. Superoxide anions are fairly unreactive whereas H_2_O_2_ and the hydroxyl radical intermediates formed during detoxification are highly reactive [179]. H_2_O_2_ is converted to a hydroxyl radical which has been found to react with and damage most metabolites and macromolecules within the cell, often producing additional radicals that lead to further damage [179]. Examples of the harmful effects of ROS interaction with proteins include loss of protein function, cross-linking, and fragmentation causing increased levels of proteolysis. Additionally, hydroxyl radicals are one of the only ROS reactive enough to interact directly with nucleic acids in DNA, which has been shown to lead to mutagenesis, carcinogenesis, and apoptosis.

Once ROS have oxidized GSH to GSSG, they can no longer react with other cellular macromolecules and GSH’s anti-oxidative properties are frozen [180]. GSH is produced in cells via the GSH-synthesis pathway that stems directly from the methyl cycle, where the *SAM1* and *SAM2* genes play essential roles, via the transsulfuration pathway [181] (Fig 9). When AdoMet is used as a methyl donor it is converted to S-AdenosylHomocysteine (AdoHcy) which feeds into production of homocysteine (Hcy) and eventually GSH. Additionally, GSH can be made available to the cell via reduction of GSSG back to GSH by glutathione reductase (*GLR1*) and NADPH [180]. GSH can also be made available to the cell through γ-glutamyltransferase (*ECM38*) and Cys-Gly metallodipeptidase (*DUG1*) via recycling cysteine from conjugated glutathione [180,182] (Fig 9).

Other important enzymes that are utilized in GSH biosynthesis include gamma-glutamyltranspeptidase (γ-GT), encoded by *ECM38*, which is responsible for degrading GSH. γ-GT is the only known enzyme that will degrade GSH to regulate GSH homeostasis [183]. γ-GT hydrolyzes GSH to form glutamate and cysteinylglycine, which can further be recycled and used in amino acid synthesis or DNA repair. *DUG1* encodes for enzyme Cys-Gly metallo-di-peptidase, which catabolizes cysteinylglycine with help from glutamine amidotransferase (encoded by *DUG2* and *DUG3*) [184]. Cys-Gly metallo-di-peptidase hydrolyzes L-cysteinyl-glycine into cysteine and glycine. Another enzyme, glutathione oxidoreductase, encoded by *GLR1,* is responsible for converting GSSG to GSH. GSH is the active state of glutathione, and is the state necessary to carryout detoxification of oxidants [185]. Glutathione oxidoreductase uses NADPH to reduce glutathione disulfide (GSSG) into two GSH molecules in their active (reduced) form. Alternatively, the enzyme phospholipid hydroperoxide glutathione peroxidase, encoded by *GPX1, GPX2,* and *HYR1,* is responsible for detoxifying harmful compounds, such as hydrogen peroxide [186,187]. They do so by oxidizing two active GSH molecules to glutathione disulfide and reducing hydrogen peroxide to two water molecules.

In *sam211/sam211* mutant cells we saw differential expression of: *MET6* (fold change of 0.55, p-value = 0.04005), *SAM1* (fold change of 0.34, p-value = 0.0002), *SAM2* presented as absent due to its complete deletion, *SAH1* (fold change of 0.46, p-value = 0.0036), *CYS3* (fold change of 0.42, p-value = 0.00005), *CYS4* (fold change of 0.50, p-value = 0.0055), *STR3* (fold change of 1.57, p-value = 0.03755)*, STR2* (fold change of 2.33, p-value = 0.0002)*, GSH1* (fold change of 0.51, p-value = 0.00145)*, GPX2* (fold change of 0.47, p-value = 0.0014)*, GRX4* (fold change of 2.52, p-value=0.00005), *GTO1* (fold change of 1.74, p-value=0.00965), and *DUG1* (fold change of 1.81, p-value = 0.00815). We did not observe any DEGs in these pathways in the *sam111/sam111* mutant cell beyond *SAM1* itself.

As previously mentioned, sodium selenite spontaneously lowers GSH concentrations through reacting with it, altering the cell’s ability to combat ROS. As seen in Fig 8A, *sam211/sam211* are sensitive to sodium selenite as compared to wildtype and *sam111/sam111* mutant cells. Reduced expression of *CYS4*, *CYS3*, and *GSH1* and increased expression of *STR3* and *STR2* (Fig 9) would point to less production of glutathione overall, such that when the cells are introduced to sodium selenite, they are more sensitive to the harmful effects due to a reduction of the neutralization mechanism. Contrastingly, *sam111/sam111* mutant cells grew equivalently as wildtype cells (Fig 8A), indicating this pathway is not increased in efficiency due to increased AdoMet. Methyl viologen, another drug that has capabilities to induce ROS, does so through production of superoxide radicals. Specifically MV donates electrons to the electron transport chain, ultimately becoming a MV cation radical, which then reacts with oxygen to form superoxide radical [188]. Although MV can raise ROS levels, the *sam211/sam211* mutant cells did not exhibit any significant growth curve differences in comparison to wildtype. The *sam111/sam111* mutant cells, on the other hand, showed only very little growth around 60 hours. Thiourea has the ability to react with and deplete GSH and NADPH, hindering the cell’s abilities to combat ROS. The *sam111/sam111* mutant cells did not exhibit any DEGs in the glutathione biosynthesis pathway (Fig 9), therefore to determine a cause for their decreased growth in MV and thiourea, further future investigation is warranted.

Sodium thiosulfate donates electrons to radical ROS species, therefore neutralizing them. In this condition, the *sam211/sam211* mutant cells exhibited an increased growth efficiency. This may be explained by an increased expression of *GRX4* (fold change of 2.52, p-value=0.00005), which is responsible for eliminating ROS species by coupling its reduction with the oxidation of GSH to GSSG along with the reduction of a glutaredoxin (Fig 9). The *sam211/sam211* mutant cells also express upregulated *GTO1* (fold change of 1.74, p-value=0.00965), part of the family of Gto proteins (composed of Gto1, Gto2, and Gto3), which is a GSH transferase, also responsible for neutralizing ROS species, specifically those that are classified as GS-hydroquinones [189,190]. Therefore, whereas in conditions of increased ROS, these mutants do not appear to sufficient glutathione to survive as well as wildtype cells, in this condition where ROS are minimized, the *sam211/sam211* cells appear to be able to combat this reduced load more efficiently through their excess of Grx4 and Gto1.

#### D. Lack of *sam2* confers resistance to arginine biosynthesis inhibitors

We identified four different conditions with impacts on arginine biosynthesis and metabolism (Supplementary Table 5). These conditions include L-glutamic acid γ-monohydroxamate (PM22D A1-4), arginine hydroxamate (PM 22D B1-4), ammonium sulfate (PM23A A9-12), and urea 2%-7% (PM9 E7-E12). The reported mechanisms of action were classified by BiOLOG as follows: L-glutamic acid γ-monohydroxamate-“tRNA synthetase”, arginine hydroxamate-“tRNA synthetase”, ammonium sulfate-“toxic cation”, and urea 2%-7%-“osmotic sensitivity”. We observed phenotypic growth differences in our *sam2Δ/sam2Δ* strains in all of these conditions. Fig 10 shows the growth curves of the wildtype, *sam111/sam111*, and *sam2Δ/sam2Δ* strains in response to these compounds in the PM plates. Multiple concentrations were tested in different PM wells and data from a representative well is shown. (For all growth curves see Supplementary Tables 6 and 7).

**Fig 10.**
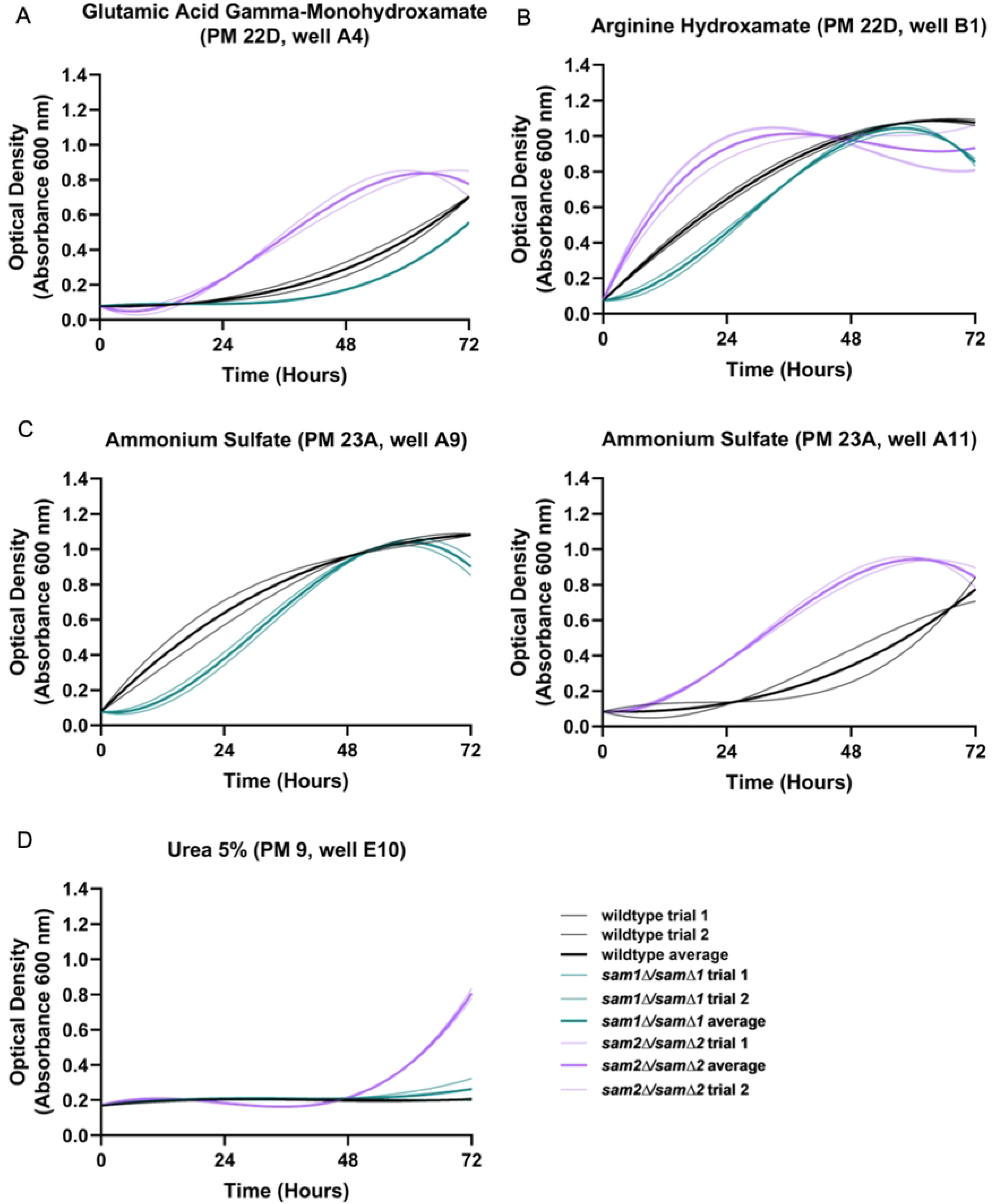
Growth curves of wildtype, *sam1Δ/sam1Δ* and *sam2Δ/sam2Δ* cells in the presence of (A) L-Glutamic Acid Gamma-Monohydroxamate, (B) Arginine Hydroxamate, (C) Ammonium Sulfate, and (D) Urea 5%. Growth curves generated from timepoint data of the Phenotypic Microarray experiments. The independent trials for each strain are shown in a lighter weight line, while the average is shown in the same color in bold. (A) Growth curves are shown for wildtype, *sam1Δ/sam1Δ, sam2Δ/sam2Δ* cells in the highest concentration of L-Glutamic Acid Gamma-Monohydroxamate (A4). (B) Growth curves are shown for wildtype, *sam1Δ/sam1Δ, sam2Δ/sam2Δ* cells in the lowest concentration of Arginine Hydroxamate (B1). (C) Growth curves are shown for wildtype and *sam1Δ/sam1Δv*cells in the lowest concentration of Ammonium Sulfate (A9) and wildtype and *sam2Δ/sam2Δ* cells in the second highest concentration (A11). (D) Growth curves are shown for wildtype, *sam1Δ/sam1Δ, sam2Δ/sam2Δ* cells in 5% Urea (E10).

In general, *sam2Δ/sam2Δ* strains exhibited increased growth compared to both wildtype and *sam1Δ/samΔ1* strains in these conditions. In PM plate 22D well A4 containing L-glutamic acid γ-monohydroxamate, (Glu-HXM) the wildtype cells began growth at 24 hours, whereas the *sam1Δ/samΔ1* mutant cells exhibited increased adaptation time with growth beginning around the 40-hr timepoint and ended with an overall lower growth efficiency compared to wildtype (Fig 10A). At the same concentration of Glu-HXM, the *sam2Δ/sam2Δ* mutant cells started to grow before the 24-hr timepoint, grew fastest between 24 and 48 hours and ended with a slightly higher efficiency to wildtype (Fig 10A). In PM plate 22D well B1 containing arginine hydroxamate and PM plate 23A well A9 containing ammonium sulfate, the *sam1Δ/samΔ1* mutant cells exhibited an increased adaptation time and slightly decreased growth rate in comparison to the wildtype (Fig 10B & 10C). In the same concentration of arginine hydroxamate, the *sam2Δ/sam2Δ* mutant cells exhibited an increased growth rate and efficiency between 0 and 24 hours in comparison to wildtype but had a similar growth efficiency by 48 hours (Fig 10B). In ammonium sulfate in PM plate 23A well A11, *sam2Δ/sam2Δ* mutant cells exhibited an initial increased growth rate and decreased adaptation time compared to wildtype, but a similar ending growth efficiency (Fig 10C). In PM plate 9 well E10 containing 5% urea, the *sam2Δ/sam2Δ* mutant cells began growth at 48 hours and maintained growth until 72 hours, whereas the wildtype cells and the *sam1Δ/samΔ1* mutant cells exhibited no significant growth (Fig 10D).

L-glutamic acid γ-monohydroxamate (Glu-HXM) and arginine hydroxamate are inhibitors of arginine metabolism, while ammonium sulfate and urea are component parts of the pathway (Fig 11). Arginine metabolism is crucial for cellular growth, stress response, metabolic regulation, nitrogen source supplementation, and polyamine synthesis [191]. Perhaps the most direct connection of these to our *SAM* genes of interest is the involvement in polyamine synthesis which is also linked to the methyl cycle and AdoMet availability. Ornithine, a compound of arginine biosynthesis and the urea cycle (Fig 11), is a precursor to putrescine, an essential precursor to polyamines; spermine and spermidine. (Lee et al. 2019). Synthesis of polyamines allows for increased DNA repair, stability, and resistance to genotoxic and oxidative damaging substances [193]. Glu-HXM is a toxic analog to glutamic acid, and therefore has the ability to inhibit synthesis of some amino acids (such as arginine and glutamate) and limit cell growth and replication [194]. Arginine hydroxamate is a toxic analog to arginine, which may promote inhibition of the TCA cycle and ATP generation through a negative feedback loop, halting arginine synthesis and therefore fumarate production by Arg4 (Fig 11) [195]. Ammonium sulfate is a source of ammonium for the cell [196]. Cation ammonium is transported in the cell by ammonium permeases, and is used as a nitrogen source within the cell [197]. Urea, on the other hand, is produced as a byproduct in the arginine biosynthesis pathway, and can be degraded to ammonium to be used as a nitrogen source [198]. In lower concentrations, urea can be beneficial to the cell providing an extra nitrogen source, but in larger concentrations, urea can form lethal ethyl carbamate, which is genotoxic to cells [199].

**Fig 11.**
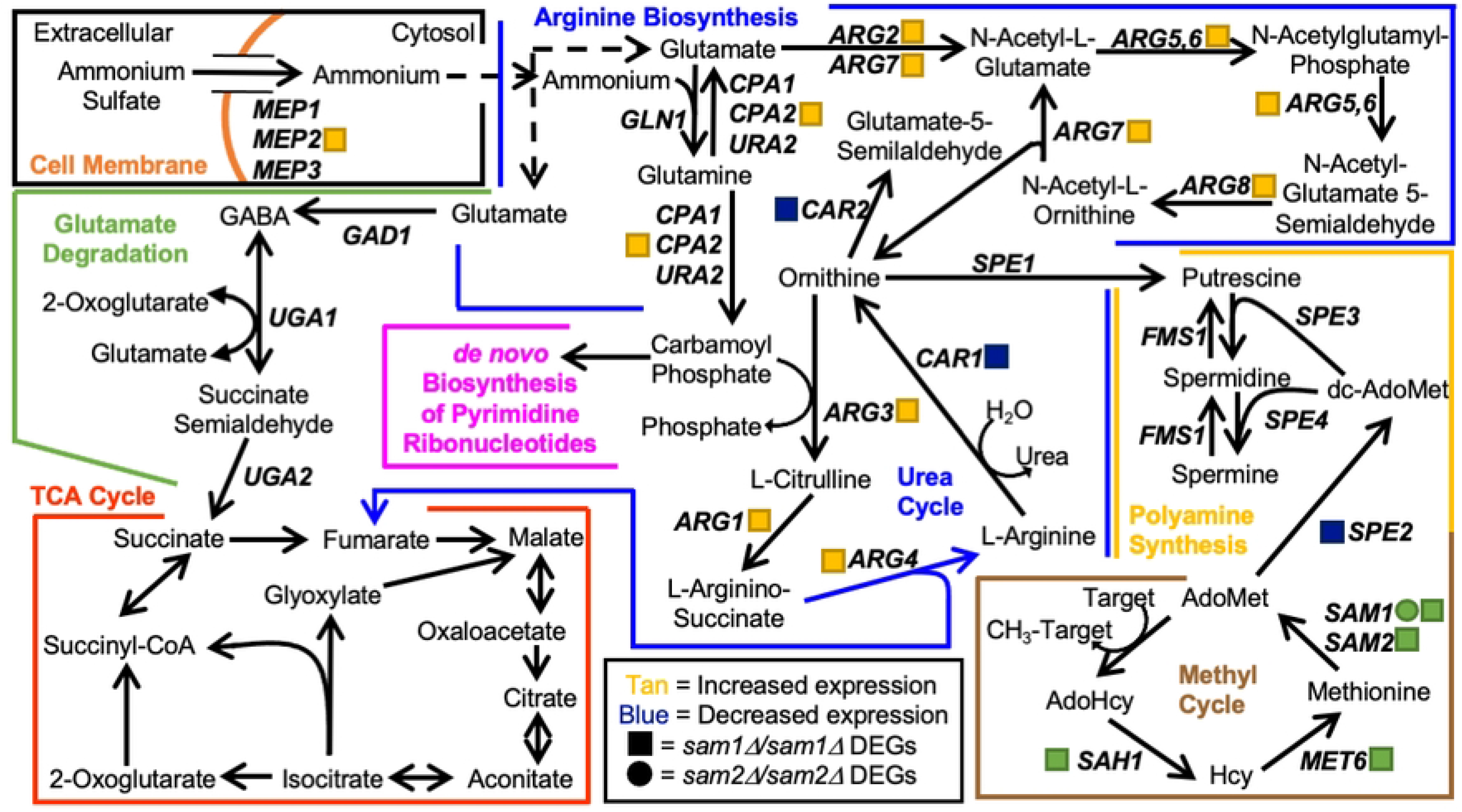
Pathways Involved in Arginine Biosynthesis. Production of arginine in *S. cerevisiae* begins with synthesis of ornithine from glutamate, then conversion to arginine in the urea cycle. In the process, several other substrates are formed that can be used in other biochemical pathways, such as the TCA cycle, polyamine synthesis, and *de novo* biosynthesis of pyrimidine ribonucleotides. The gene names that encode the enzymes that can function in compound conversion in the pathway are given by their standard names. *sam2Δ/sam2Δ* DEGs are represented by red (increased expression) and green (decreased expression) squares. sam1Δ/sam1Δ DEGs are represented by red (increased expression) and green (decreased expression) circles.

Through investigation of the full list of DEGs from the *sam1Δ/samΔ1* and *sam2*11*/sam2*11 mutant strains, we were able to identify *MEP2, CPA2, ARG2, ARG7, ARG5,6, ARG8, ARG3, ARG1, ARG4,* and *CAR1* as genes involved in arginine metabolism, and *MET6, SAH1, SAM1, SAM2,* and *SPE2* involved in the methyl cycle and polyamine synthesis. RNA-sequencing did not reveal any differentially expressed genes within this pathway between *sam1Δ/sam1Δ* and wildtype, except for no expression of *SAM1* due to the deletion of the gene. We explored differentially expressed genes in our *sam2*11*/sam2*11 mutant within these pathways to determine if the observed phenotypic growth differences could be explained at this level of regulation.

Interestingly, almost every gene in the arginine biosynthesis pathway is upregulated in *sam2*11*/sam2*11 mutant cells. This includes: *CPA2* (fold change of 3.05, p-value=0.00005), *ARG2* (fold change of 1.57, p-value=0.0345), *ARG7* (fold change of 3.00, p-value=0.00005), *ARG5,6* (fold change of 2.38, p-value=0.0005), *ARG8* (fold change of 2.02, p-value=0.00125), *ARG3* (fold change of 4.89, p-value=0.00005), *ARG1* (fold change of 2.51, p-value=0.00045), and *ARG4* (fold change of 1.98, p-value=0.00995). With these DEGs differentially overexpressed, *sam2*11*/sam2*11 mutant cells may not only produce more of the amino acid arginine, but also supply more fumarate to the TCA cycle via Arg4 (Fig 11). Increased expression of *CPA2* may allow for increased synthesis of substrate carbamoyl phosphate, not only to be used in the urea cycle subsection of arginine biosynthesis, but also for *de novo* biosynthesis of pyrimidine ribonucleotides (which is discussed in section E).

Arginine hydroxamate, a known toxic analog to arginine, inhibits arginine biosynthesis via negative feedback [195]. Previous studies, however, have found that increased concentrations of ornithine and citrulline are able to reverse the inhibitory effect of arginine hydroxamate [195], both of which are produced in the urea cycle, a subset of the arginine metabolism pathway (Fig 11). We hypothesize that because the *sam2*11*/sam2*11 mutant cells have upregulation of *ARG3* and *ARG7,* these cells therefore have more ornithine and citrulline available to combat the inhibitory effect of arginine hydroxamate, allowing these cells to grow better in this condition. Furthermore, *sam2*11*/sam2*11 mutant cells may have excess arginine via upregulated Arg4, allowing these mutants to be less susceptible to the effects as an arginine analog. The *sam1Δ/samΔ1* mutant cells had no DEGs present in the arginine metabolism pathway and grew similarly to the wildtype in this condition.

Downregulated DEGs present within the arginine metabolism pathway of *sam2*11*/sam2*11 mutant cells include *CAR1* (fold change of 0.56, p-value=0.00925), and *CAR2* (fold change of 0.29, p-value=0.00005), both of which are involved in arginine degradation. Urea is a byproduct in the urea cycle during conversion of arginine to ornithine by Car1 (Fig 11), and is commonly used as a nitrogen source [198]. Urea can be degraded to carbon dioxide and ammonium, however, in higher concentrations, it may react with ethanol to form ethyl carbamate, a toxic compound that can induce genotoxicity [199,200]. Because *sam2*11*/sam2*11 mutant cells downregulate *CAR1*, they may be able to hinder the effects of ethyl carbamate production in the presence of excess urea, as the downregulation likely created a lower urea concentration as their starting point (Fig 11). We hypothesize this contributed to the *sam2Δ/sam2Δ* mutant cells growth in excess urea, when the wildtype cells and the *sam1Δ/samΔ1* mutant cells (with no downregulation) were fully inhibited.

Glu-HXM is a toxic analog to glutamine, an important substrate used in arginine biosynthesis, and subsequently the TCA cycle (Fig 11). The mechanism for Glu-HXM toxicity in *S. cerevisiae* has not yet been examined, however, in *E. coli*, Glu-HXM inhibits glutamine-fructose-6-phosphate amidotransferase via competitive inhibition of the glutaminase binding site [201]. In *S. cerevisiae*, glutamine-fructose-6-phosphate amidotransferase is encoded by the gene *GFA1,* and while it is not a DEG in either mutant strain, we hypothesize that the *sam2Δ/sam2Δ* mutant cells had an increased growth rate and decreased adaptation time because the multiple upregulated DEGs throughout the arginine metabolism pathway would allow these cells to better deal with inhibition at this single point (Fig 11). Alternatively, the *sam1Δ/samΔ1* mutant cells show slightly impaired growth compared to wildtype but did not exhibit any DEGs in relation to the arginine metabolism pathway (Fig 11). Therefore, the growth difference is not likely linked to this pathway, and further research is warranted.

Ammonium is a nitrogenic cation that is important for several cellular functions, including serving as a nitrogen source for amino acid catabolism and inhibiting autophagy [202]. *S. cerevisiae* grown in media rich in ammonium sulfate, were found to uptake this compound as ammonium cations through membrane permeases encoded by genes *MEP1, MEP2,* and *MEP3* [203]. Interestingly, *sam2*11*/sam2*11 mutant cells contain upregulated *MEP2* (fold change of 2.54, p-value=0.0061). Ammonium is not only used as a nitrogen source, but also participates as a substrate for the synthesis of glutamine from glutamate (Fig 11) and is therefore essential in glutamine and arginine biosynthesis. We hypothesize that because *sam2*11*/sam2*11 mutant cells have upregulated *MEP2*, they can uptake more ammonium than wildtype and *sam1Δ/sam1Δ* mutant cells, which provides more substrate for glutamine synthesis, and therefore arginine metabolism, TCA cycle and ATP production, and polyamine synthesis. These together could contribute to the increased growth in the PM plate that the *sam2*11*/sam2*11 mutant cells exhibited.

#### E. Lack of *sam2* confers resistance to DNA synthesis inhibitors

The PM plates contain seven different conditions that could have impacts on DNA synthesis or the folate cycle (Supplementary Table 5). These conditions include 5-fluorodeoxyuridine (PM25D D9-12), hydroxylamine (PM23A C5-8), hydroxyurea (PM25D A1-4), 5-fluorocytosine (PM25D G1-4), azaserine (PM22D F1-4), 5-fluorouracil (PM25D H9-12), and 6-azauracil (PM24C D1-D4). The reported mechanisms of action were classified by BiOLOG as follows: 5-fluorodeoxyuridine - “DNA synthesis inhibitor”, hydroxylamine - “DNA damage, mutagen, antifolate (inhibits thymine and methionine synthesis)”, hydroxyurea - “Ribonucleotide DP reductase inhibitor, antifolate (inhibits thymine and methionine synthesis)”, 5-fluorocytosine - “DNA synthesis inhibitor”, azaserine-“nucleic acid inhibitor, purine, glutamine analog”, 5-fluorouracil - “nucleic acid analog, pyrimidine”, and 6-azauracil-“nucleic acid analog, pyrimidine”. We observed phenotypic growth differences in our *sam2Δ/sam2Δ* strains in four of these conditions: 5-fluorodeoxyuridine, 5-fluorocytosine, 6-azauracil, and hydroxylamine. The broad mechanism of action of these drugs are as DNA synthesis inhibitors. Fig 12 shows the growth curves of the wildtype, *sam111/sam111*, and *sam2Δ/sam2Δ* strains in response to these compounds in the PM plates. Multiple concentrations were tested in different PM wells and data from a representative well is shown. (For all growth curves see Supplementary Tables 6 and 7).

**Fig 12.**
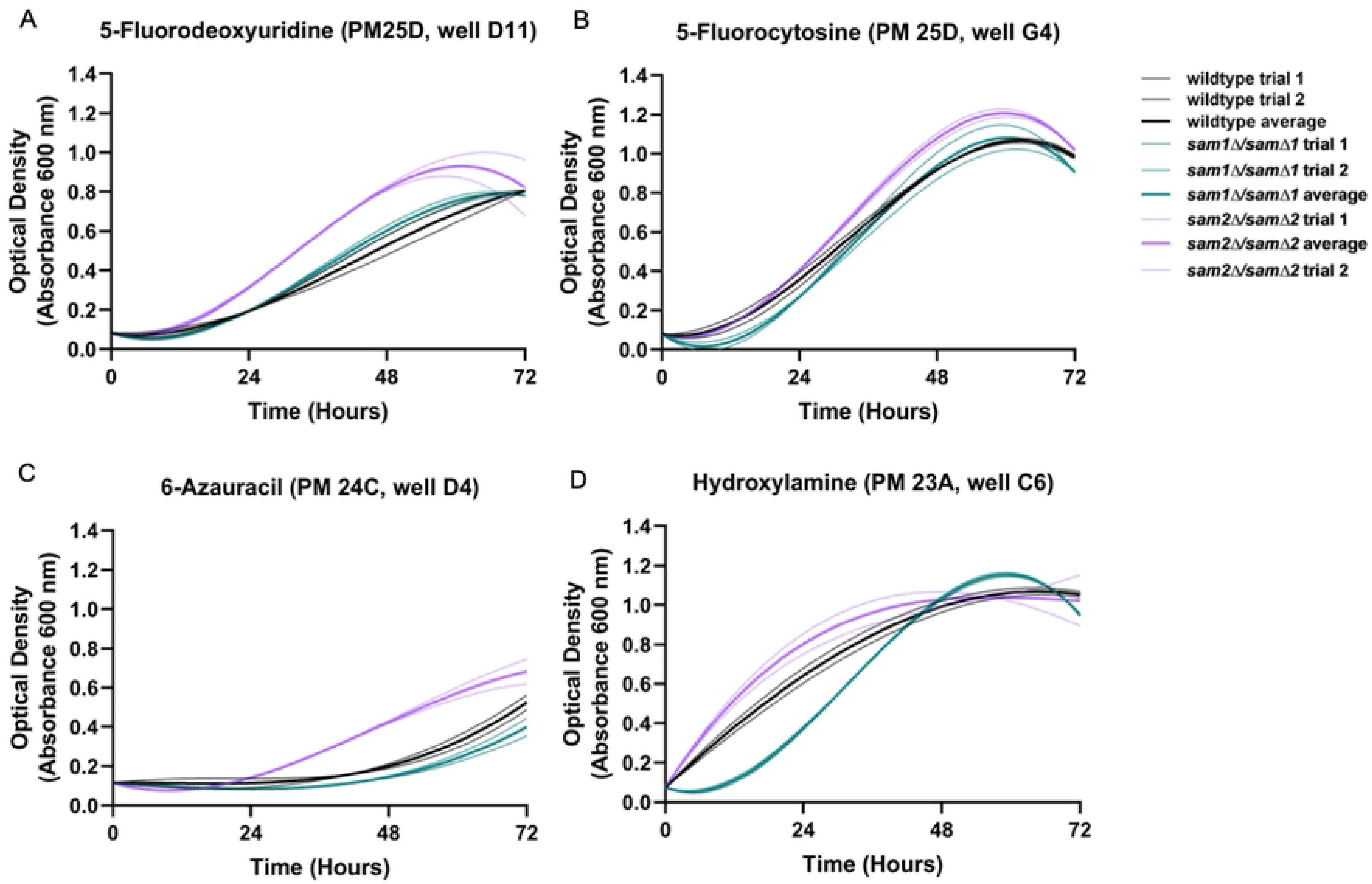
Growth curves of wildtype, *sam1Δ/sam1Δ* and *sam2Δ/sam2Δ* cells in the presence of (A) 5-Fluorodeoxyuridine, (B) 5-Fluorocytosine, (C) 6-Azauracil, and (D) Hydroxylamine. Growth curves generated from timepoint data of the Phenotypic Microarray experiments. The independent trials for each strain are shown in a lighter weight line, while the average is shown in the same color in bold. (A) Growth curves are shown for wildtype, *sam1Δ/sam1Δ, sam2Δ/sam2Δ* cells in the second highest concentration of 5-Fluorodeoxyuridine (D11). (B) Growth curves are shown for wildtype, *sam1Δ/sam1Δ, sam2Δ/sam2Δ* cells in the highest concentration of 5-Fluorocytosine (G4). B) Growth curves are shown for wildtype, *sam1Δ/sam1Δ, sam2Δ/sam2Δ* cells in the highest concentration of 6-Azauracil (D4). B) Growth curves are shown for wildtype, *sam1Δ/sam1Δ, sam2Δ/sam2Δ* cells in the third highest concentration of Hydroxylamine (C6).

In all conditions presented here, the wildtype and *sam2Δ/samΔ2* mutant cells began to grow between the 0 and 24-hr timepoint and maintained increased growth through 48 or 72 hours. Generally, *sam2Δ/sam2Δ* strains exhibited increased growth compared to both wildtype and *sam1Δ/samΔ1* strains in these conditions. In PM plate 25D well D11 containing 5-fluorodeoxyuridine, the wildtype and *sam1Δ/samΔ1* mutant cells began to grow around the 12-hr timepoint and maintained increased growth throughout the remainder of the experiment (Fig 12A), with no significant difference in the growth patterns between the two. At the same concentration of 5-fluorodeoxyuridine, the *sam2Δ/sam2Δ* mutant cells exhibited decreased adaptation time, with growth beginning at the 12-hr timepoint, and increased growth rate between 24 and 48 hours (Fig 12A). In PM plate 25D well G4 containing 5-fluorocytosine, the wildtype and *sam1Δ/samΔ1* mutant cells grew similarly, beginning to grow before the 24-hour timepoint and starting to level off after 48-hours. In contrast, the *sam2Δ/sam2Δ* mutant cells exhibited increased growth rate and efficiency (Fig 12B). In PM plate 24C well D4 containing 6-azauracil, growth for both the wildtype and *sam1Δ/samΔ1* mutant cells began between the 24-hour and 48-hour timepoints, and they maintained increased growth for the rest of the experiment. In the same concentration of 6-azauracil, *sam2Δ/sam2Δ* mutant cells began to grow at the 24-hour time point with decreased adaptation time, and continued growth until the end of the experiment, exhibiting increased growth rate, and increased efficiency of growth compared to wildtype (Fig 12C). In PM plate 23A well C6 containing hydroxylamine, the *sam1Δ/samΔ1* mutant cells exhibited decreased growth rate and an increased adaptation time with growth beginning around 12 hours, in comparison to wildtype cells, (Fig 12D). In the same concentration of hydroxylamine, the *sam2Δ/sam2Δ* mutant cells exhibited a slight decrease in adaptation time but otherwise similar growth compared to wildtype (Fig 12D).

5-fluorodeoxyuridine, 5-fluorocytosine, 6-azauracil, and hydroxylamine all inhibit DNA synthesis through altering deoxyribonucleoside triphosphate (dNTP) levels. As the building blocks of DNA, dNTPs are utilized in replication and repair processes. dNTP pools are controlled by two major pathways: the *de novo* biosynthesis pathway and the salvage pathway. dNTP levels within the cell must be highly regulated in order to maintain genomic stability [204]. This balance is primarily accomplished via the ribonucleotide reductase (RNR) complex in *de novo* synthesis (Fig 13). Nucleotide salvage pathways, on the other hand, are primarily regulated though feedback inhibition and gene regulation [205]. The cell balances these two regulation mechanisms based on specific cellular conditions, where nucleotide salvage is preferred over *de novo* synthesis when substrate is available [206]. Studies have found that the nucleotide retrieval source may determine where it will be used moving forward. For example, nucleotides that were synthesized from the *de novo* pathway are likely to be used in incorporation into UDP sugars, and nucleotides that were salvaged from pyrimidines or purines are likely to be used in future synthesis of RNA [207]. RNR can catalyze the conversion of ribonucleotides to deoxyribonucleotides [207]. The RNR complex is the rate-determining step in *de novo* biosynthesis of pyrimidines and purines, and is regulated in multiple ways: transcriptional regulation of *RNR3* following DNA damage, allosteric activation or inhibition via ATP or dATP binding, and via a small protein called Sml1 in response to environmental stressors or genetic damage [208–211]. Alternatively, nucleotide salvage pathways are the series of catabolic processes in which a nutrient-deprived cell may recover essential bases or nucleosides from exogenous sources or following DNA/RNA degradation [212]. The pyrimidine salvage pathway allows for the catabolism of 2-deoxycytidine to form pyrimidine bases uracil, thymine, and cytosine. The salvage and *de novo* pathways interconnect with the folate cycle, and ultimately the methyl cycle, through shared intermediates. These linked pathways were explored to determine if expression differences might explain the observed consistent increased survival of *sam2Δ/sam2Δ* mutant cells in the presence of 5-fluorodeoxyuridine, 5-fluorocytosine, and 6-azauracil.

**Fig 13.**
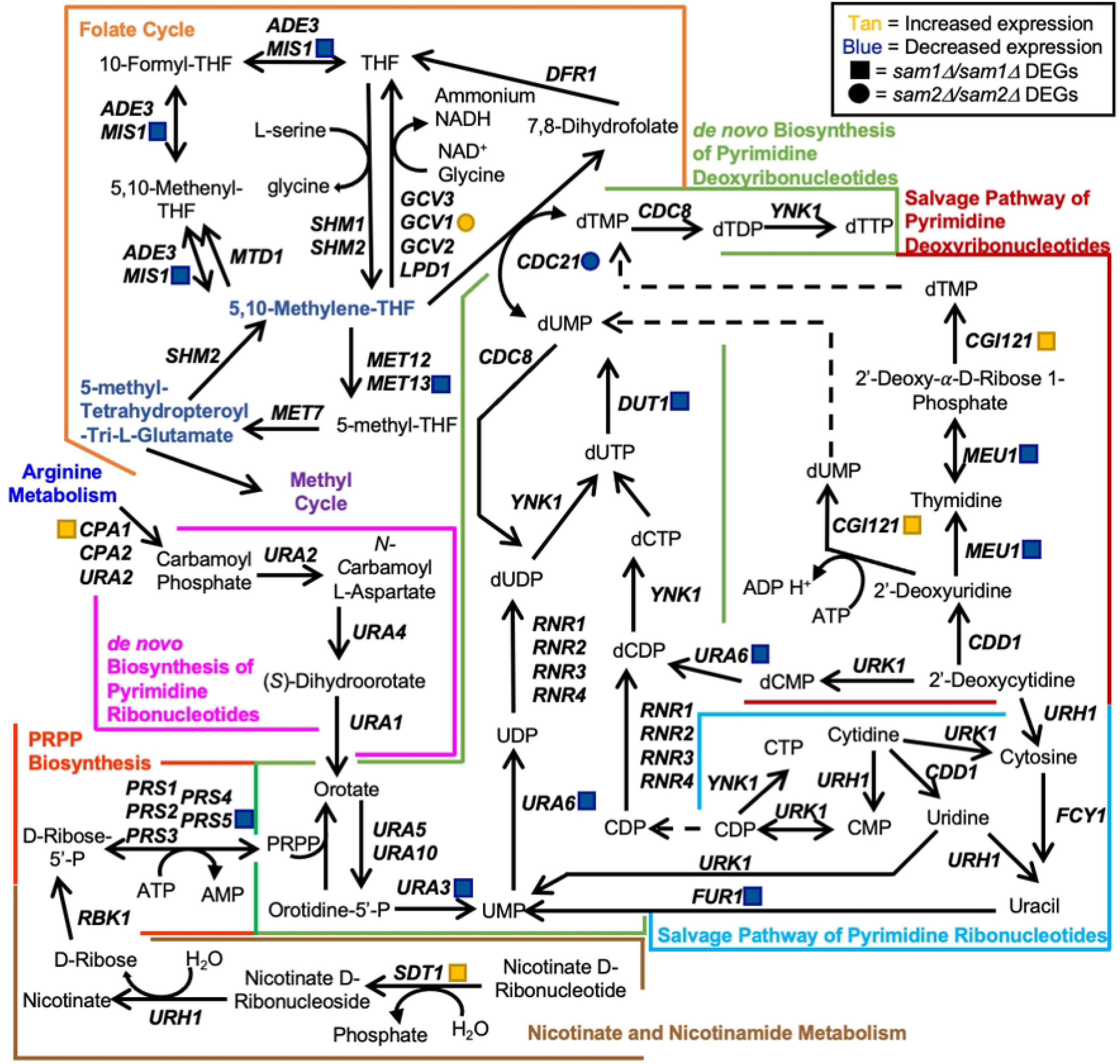
The Interconnection between the Folate Cycle, *de novo* Biosynthesis of Pyrimidine Nucleotide Pathways, Salvage Pathway of Pyrimidine Nucleotides, Phosphoribosyl Diphosphate (PRPP) biosynthesis, and Nicotinate and Nicotinamide Metabolism. dNTP pools are maintained through several pathways, including the *de novo* and salvage pathways of pyrimidines, PRPP biosynthesis, nicotinate and nicotinamide metabolism, and the folate cycle. The gene names that encode the enzymes that can function in compound conversion in the pathway are given by their standard names. *sam2Δ/sam2Δ* DEGs are represented by red (increased expression) and green (decreased expression) squares. *sam1Δ/sam1Δ* DEGs are represented by red (increased expression) and green (decreased expression) circles. The conversion of dCTP to dUTP is catalyzed by a dCTP deaminase, though the gene encoding this enzyme has not been identified and thus is not labeled in the figure. Through research on homologous genes, we believe that the dCTP deaminase in *S. cerevisiae* could be DCD1 [218,219]. Though this is not confirmed, and DCD1 is not a DEG, and thus not included.

5-fluorodeoxyuridine, 5-fluorocytosine, 6-azauracil, and hydroxylamine all impact the pyrimidine deoxyribonucleotide *de novo* biosynthesis pathway, leading to similar cellular impacts. These drugs, however, exert their effects at varying points in the pathway and act in different manners. 5-Fluorodeoxyuridine irreversibly binds thymidylate synthase (TS, referred to as TS/Cdc21 moving forward), an essential enzyme found in the *de novo* biosynthesis of pyrimidine nucleotides pathway (Fig 13). TS/Cdc21 catalyzes the conversion of deoxyuridine monophosphate (dUMP) to deoxythymidine monophosphate (dTMP). When TS/Cdc21 activity is inhibited by 5-fluorodeoxyuridine, deoxythymidine triphosphate (dTTP) becomes depleted, and DNA synthesis is inhibited. At high concentrations, 5-fluorodeoxyuridine can lead to a “thymineless death” through inhibition of DNA replication due to loss of dTTP [213]. Other consequences of TS/Cdc21 inhibition leading to cell death have been proposed, such as imbalances in pyrimidine nucleotides, and ineffective DNA repair pathways with toxic intermediates. Similar to 5-flurodeoxyuridine, 5-fluorocytosine (5-FC) interrupts DNA synthesis by targeting TS/Cdc21, but also has additional roles in impairing protein synthesis. Nontoxic 5-fluorocytosine must first be converted into toxic 5-fluorouracil (5-FU) by cytosine deaminase encoded by *FCY1*. Kinetic studies have shown that the rate-limiting step in activating 5-fluorocytosine into its toxic form is the release of 5-FU from Fcy1 [214]. 5-FU is then converted by the product of *FUR1*, uracil phosphoribosyltransferase, to toxic 5-fluorodeoxyuridine 5’-monophosphate (5FdUMP) and 5-fluorouridine 5’-monophosphate (5FUMP). These compounds lead to the disruption of both DNA and protein synthesis; 5FdUMP irreversibly inhibits TS/Cdc21, consequently inhibiting DNA synthesis, while 5FUMP inhibits protein synthesis through incorporation of its metabolites into mRNA, tRNA, and rRNA. 5FdUMP exerts its toxic effects by forming a complex with TS/Cdc21 and 5,10-methylene-THF. (Interestingly, despite 5-FC conversion to 5-FU in the cell, our cells grown directly in the presence of 5-FU did not show altered growth. This could speak to limitations of detecting growth impacts with only the four concentrations present. Concentration data is proprietary to BiOLOG so we are unable to make direct comparisons on the 5-FC and 5-FU concentrations tested.) The uracil analog, 6-azauracil (6-AU) also inhibits DNA synthesis through its impact on the purine and pyrimidine pathways. 6-AU is known to deplete intracellular GTP and UTP concentrations through inhibition of IMP dehydrogenase and orotidylate carboxylase, respectively. Orotidylate carboxylase, encoded by *URA3,* is an enzyme that is necessary for the *de novo* biosynthesis of pyrimidines converting orotidine-5’-P to UMP (Fig 13) [215]. Disruption at this point in the *de novo* pyrimidine biosynthesis pathway has been shown to significantly reduce cellular nucleotide levels and inhibit cellular growth [216]. Hydroxylamine (HA) has previously been identified in bacteria to be an RNR complex inhibitor when methylated [217]. Because RNR is the rate-limiting step in *de novo* pyrimidine biosynthesis, inhibition could create a deoxyribonucleotide pool imbalance.

Through investigation of the full list of DEGs from the *sam1Δ/samΔ1* and *sam2*11*/sam2*11 mutant strains, we were able to identify *URA3, URA6, DUT1, and CDC21,* of the *de novo* biosynthesis of pyrimidine nucleotides pathway; *CGI121*, *MEU1,* and *FUR1* of the pyrimidine salvage pathways; *MIS1, GCV1,* and *MET13* of the folate cycle; *PRS5* of the PRPP biosynthesis; *CPA2* of arginine metabolism, and *SDT1* of nicotinate and nicotinamide metabolism as showing altered expression. We explored differentially expressed genes in our mutant strains within these pathways to determine if the observed phenotypic growth differences could be explained at this level of regulation.

In *sam2Δ/sam2Δ* mutant strains, genes in the pyrimidine dNTP salvage pathway were found to have altered expression. Increased expression (fold change of 1.74, p-value=0.00655) of *CGI121* and decreased expression (fold change of 0.55, p-value=0.00815) of *MEU1* would appear to enable the cell to direct 2’-deoxyuridine towards conversion to dUMP and 2’-deoxy-alpha-D-ribose 1-phosphate toward conversion to dTMP, and away from thymidine (Fig 13). Furthermore, we have shown *sam2Δ/sam2Δ* mutant cells have decreased amounts of AdoMet [2]. Because ATP is a component in the conversion of methionine to AdoMet, and both *SAM1* and *SAM2* are downregulated in the *sam2Δ/sam2Δ* mutant strain, there could potentially be more ATP available to other reactions in these cells. The conversion of 2’-deoxyuridine into dUMP by Cgi121 may produce more dUMP, with ATP more readily available and *CGI121* upregulated. Any potential increases in dTMP through increased expression of *CGI121*, might also assist in counteracting the depletion of dTTP and account for the *sam2Δ/sam2Δ* strains showing resistance, as compared to wildtype, to the 5-fluorodeoxyuridine and 5-FC conditions that target this portion of the pathway. As well, increased levels of dUMP would be available to the reaction that TS/Cdc21 catalyzes in the *de novo* biosynthesis of pyrimidine nucleotides pathway. Because TS/Cdc21 is targeted by 5-fluorodeoxyuridine, increased dUMP substrate being available for TS/Cdc21 could help negate these inhibitory effects. Ultimately, gene expression increases in the pyrimidine salvage pathway could help avoid the thymineless death under 5-fluorodeoxyuridine providing needed substrates through altered pathway usage. 5-FC also inhibits DNA synthesis like 5-fluorodeoxyuridine and 5-fluorouracil, however, it must be processed in additional steps prior to TS/Cdc21 inhibition. Initially 5-FC is converted to 5-FU by cytosine deaminase (Fcy1) within the salvage pathway of pyrimidine ribonucleotides, which can bind this enzyme due to its similar structure to cytosine [220]. 5-FU, an uracil analog, is then converted by uracil phosphoribosyl transferase (Fur1) to toxic metabolite 5FdUMP. 5FdUMP irreversibly binds to TS/Cdc21 in a ternary complex with 5,10-methylene-THF, abolishing normal function of TS/Cdc21 [221]. The binding to TS/Cdc21 inhibits dTMP production, therefore causing cytotoxicity [222]. *sam2Δ/sam2Δ* cells exhibited a significant decrease in expression of *FUR1* (fold change of 0.65, p-value 0.0321). Previous research has shown human colon tumor cells with increased expression of *UPRT* (the *FUR1* homolog) have increased sensitivity to 5-FC, as UPRT plays an essential role in converting 5-FC to its toxic forms [223]. Additionally, various mutations in *FUR1* in *S. cerevisiae* have previously been shown to confer resistance to 5-FC [223,224]. The decreased sensitivity to 5-FC in our *sam2Δ/sam2Δ* cells could therefore be explained, at least in part, by the decreased *FUR1* expression, as conversion to the toxic 5-FU would be slowed due to reduced Fur1 levels. Further research has indicated resistance to 5-FU via overexpression of genes *CPA1* and *CPA2* [225], which encode for the carbamoyl phosphate synthetase enzyme, responsible for conversion of glutamine into carbamoyl phosphate (Fig 11), which can then be used to generate pyrimidine ribonucleotides and deoxyribonucleotides (Fig 13), ultimately stimulating pyrimidine synthesis altogether. *CPA2* is an upregulated DEG in the *sam2Δ/sam2Δ* cells (fold change of 3.05, p-value=0.00005) and is active within the L-arginine metabolism pathway (Fig 13), as discussed in Section D above, and thus this additional gene expression difference may also contribute to the faster growth rate and lessened adaptation time of these cells. We hypothesize that upregulated *CPA2* would confer resistance to 5-FU, because increased production of carbamoyl phosphate would stimulate pyrimidine synthesis and potentially offset the effects of this drug.

6AU, another drug that targets the pyrimidine *de novo* biosynthesis pathway, is known to inhibit the enzyme Ura3 [226]. Interestingly, in *sam2Δ/sam2Δ* cells, there was a significant downregulation of *URA3* (fold change of 0.50, p-value = 0.0037), and subsequent genes in the *de novo* pathway: *URA6* (fold change of 0.64, p-value = 0.0354) and *DUT1* (fold change of 0.58, p-value = 0.00765). With 6-AU targeting Ura3, and the already decreased expression of *URA3, URA6*, and *DUT1*, could lead to significant decreases in dNTP levels, severely impeding the survival of these cells. *sam2Δ/sam2Δ* cells, however, had a faster adaptation time, increased rate, and increased efficiency of growth compared to wildtype. Additional research suggests, however, that sensitivity/resistance to 6AU may, in fact, not be as significantly related to expression of Ura3 [227]. Instead, the effects of 6AU have been linked to expression of pyrimidine nucleotidase, an enzyme encoded by *SDT1.* Though not directly involved in nucleotide synthesis, Sdt1 is responsible for production of important ribosides, such as active metabolites nicotinamide riboside and nicotinic acid riboside through dephosphorylation from their nucleoside 5’-monophosphate precursor molecules [227,228], and can be traced to the *de novo* pathway of pyrimidine deoxyribonucleotides through formation of substrate PRPP (Fig 13). In fact, studies suggest that there is competition between the NAD biosynthetic pathway and the nucleotide biosynthetic pathways for substrates PRPP and ATP [229]. Furthermore, overexpression of *SDT1* causes higher dephosphorylation of UMP [228]. When uridine kinase (Urk1) interacts with 6AU, it produces toxic 6-AzUMP (a phosphorylated form), which competitively inhibits Ura3 [226,230]. When 6-AzUMP is dephosphorylated, namely by Sdt1, the compound is inactivated, and therefore prevented from inhibition of Ura3, ultimately rendering 6AU inert. Through investigative phosphatase assays, it was found that Sdt1 is likely an UMP phosphatase [228]. *SDT1* is overexpressed in *sam2Δ/sam2Δ* mutant cells (fold change of 2.51, p-value = 0.00005). This large and significant overexpression could be contributing to the observed increase in growth of our *sam2Δ/sam2Δ* cells compared to wildtype in this condition. As well, it is likely that Sdt1 can confer hyposensitivity to 5-FC because of its ability to target UMP and its toxic derivatives, including 6-AzUMP and 5-FUMP, rendering them harmless [228]. In *sam1Δ/sam1Δ* mutant cells the expression of *SDT1* did not show any significant differences to that of the expression in wildtype cells, which is consistent with not observing altered growth of these cells in 6-AU or 5-FC.

Research from the 1970s found that HA reacts with the formyl group in 5-formyltetrahyrofolate, and likely other folates, in *S. cerevisiae* [231]. This reaction forms formoxime, which was shown as an inhibitor of thymidylate synthetase in *Escherichia coli* [232]. In another bacterium, *Mycobacterium bovis,* HA has been shown to target the RNR complex. As in these bacteria, *S. cerevisiae* has RNR proteins and thymidylate synthase (as discussed above). The *sam1Δ/samΔ1* mutant cells likely exhibited decreased growth in this condition due to downregulated *CDC21* expression (fold change of 0.62, p-value=0.0247), which encodes thymidylate synthase, and/or because RNR complex inhibition could create dNTP pool imbalances which in concert with less Cdc21 could cause more difficulty to interconvert between some deoxyribonucleotides in these mutant cells, thereby creating sensitivity to this condition (Fig 13).

#### F. Lack of *sam2* confers resistance to tamoxifen

The PM plates contain tamoxifen (PM24C H5-8), another known chemotherapeutic. The reported mechanism of action for tamoxifen was classified by BiOLOG as “binds estrogen receptors”. We observed phenotypic growth differences in both of our mutant strains in this condition. Fig 14 shows the growth curves of the wildtype, *sam111/sam111*, and *sam2Δ/sam2Δ* strains in response to tamoxifen in the PM plates. Multiple concentrations were tested in different PM wells and data from a representative well is shown for each mutant. (For all growth curves see Supplementary Tables 6 and 7).

**Fig 14.**
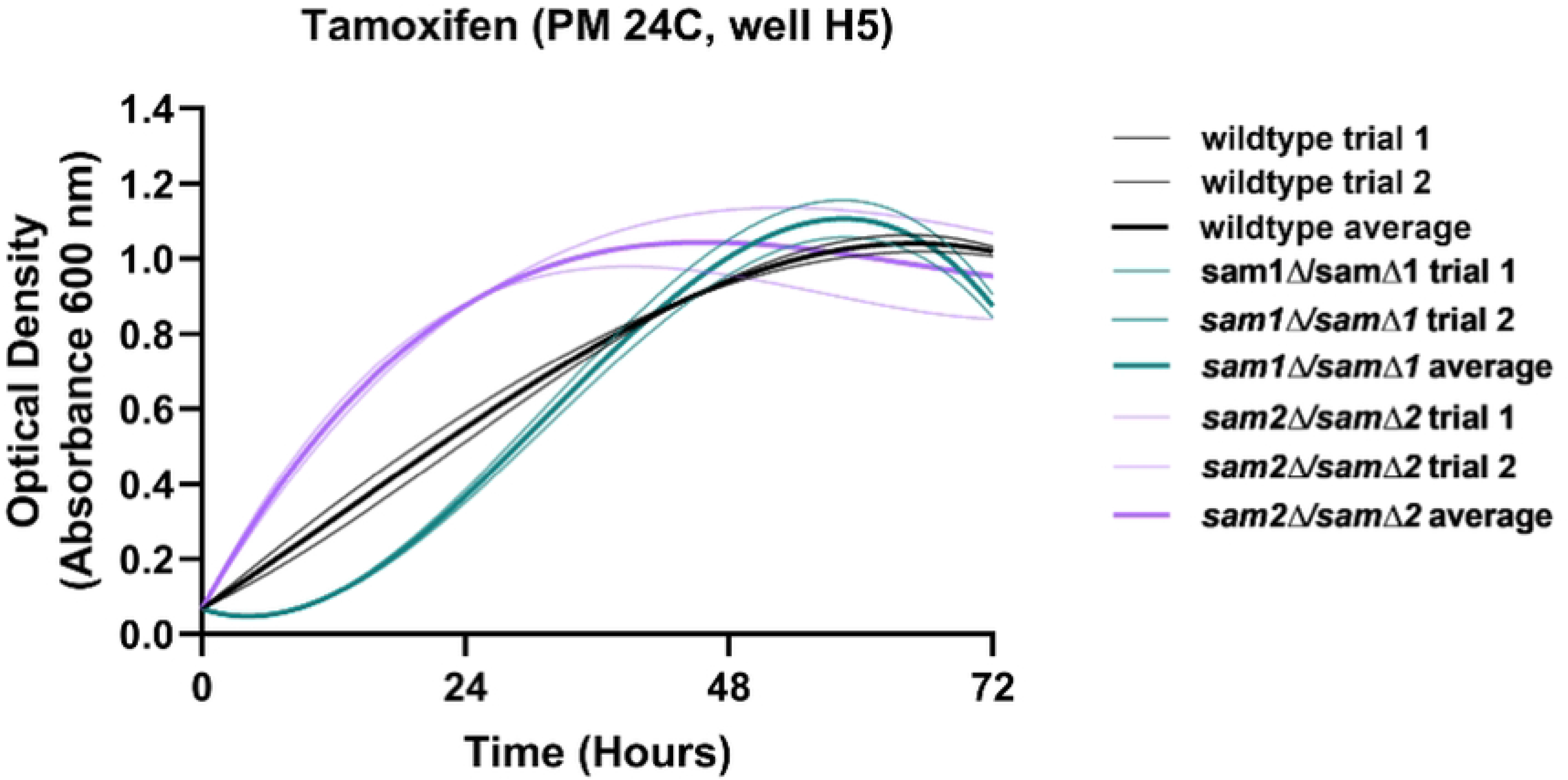
Growth curves of wildtype, *sam1Δ/sam1Δ* and *sam2Δ/sam2Δ* cells in the presence of Tamoxifen. Growth curves generated from timepoint data of the Phenotypic Microarray experiments. The independent trials for each strain are shown in a lighter weight line, while the average is shown in the same color in bold. Growth curves are shown for wildtype, *sam1Δ/sam1Δ, sam2Δ/sam2Δ* cells in the lowest concentration of Tamoxifen (H5).

PM plate 24C wells H5-8 have increasing concentrations of tamoxifen. In representative well H5 we observed that *sam2Δ/sam2Δ* mutant cells showed decreased adaptation time and an increased growth rate and efficiency to 48 hours, finalizing at a similar efficiency to wildtype at 72 hours. The *sam1Δ/sam1Δ* cells showed an increased adaptation time and slower growth rate to 12 hours, but caught up to wildtype around 40 hours. (Fig 14). Data from a representative well is shown for each mutant. (For all growth curves see Supplementary Tables 6 and 7).

Tamoxifen has previously been shown to possess antifungal activity through inhibition of calmodulin binding to calcineurin [233,234]. Calcineurin and calmodulin play crucial roles in promoting calcium ion signaling pathways in the cell, which assist in regulation of enzymatic activity, metabolic functions, and stress response [235]. Calcineurin is a protein complex made up of one calcineurin A subunit and one calcineurin B subunit. In order for calcineurin to be activated, it must first accept a calmodulin molecule bound to its calmodulin recognition site within the calcineurin regulatory domain [236]. Calmodulin, encoded by gene *CMD1,* is activated when intracellular concentrations of calcium ions increase and bind to the calcium sensor receptor, which allows for calmodulin to then bind to calcineurin, encoded by genes *CNA1* and *CMP2.* [237]. Calcium ions enter the plasma membrane through P-type ATPase pumps encoded by *PMR1*. Calcium ions can also enter the cytosol through activation of vacuolar calcium ion transporter Vcx1, an antiporter present in vacuolar membranes that pumps calcium ions into the cytosol and hydrogen ions out of the cytosol into the vacuole. Interestingly, studies have found that calcineurin inhibits the Vcx1 antiporter and instead induces the calcium ion ATPase, Pmr1, indicating that the main source of calcium ion for calmodulin is the membrane ATPase [238]. Once bound, the active calcineurin-calmodulin complex dephosphorylates target proteins, allowing for cellular responses [239]. One of these target proteins is the Crz1 transcription factor, which activates stress response genes [240]. Crz1 mediates transcription of target gene *CMK2,* a calmodulin-dependent protein kinase which phosphorylates the calcium ion channel protein Pmr1 (therefore inactivating the membrane ATPase) and vacuolar channel protein Pmc1, ultimately lowering the calcium ion concentration in the cytosol [241]. In turn, this change in abundance of calcium ion pumps and therefore intracellular calcium affects multiple cellular processes, including membrane and cytoskeleton organization, transcription of genes such as *RCH1*, a negative regulator of calcium transport in the plasma membrane, and negative feedback regulation of calmodulin [212,242]

Previous research has suggested that increased concentrations of calmodulin have the ability to suppress the effects of tamoxifen [243]. Tamoxifen induces increased intracellular calcium ion signaling through induced nuclear localization of transcription factor Crz1 followed by transcription of *CMK2*, and then inhibition of calmodulin via negative feedback [244].

The *sam2Δ/sam2Δ* mutant cells had one DEG within the calcineurin/calmodulin signaling pathway, *FAR1* (fold change of 2.05, p-value = 0.00055). *FAR1* encodes for a cyclin-dependent kinase inhibitor, responsible for inhibiting Msn5 [245]. Msn5 is responsible for nuclear import of proteins, including the Crz1 enzyme (Fig 15) [246]. When Msn5 is inhibited by Far1, Crz1 is not localized to the nucleus, and therefore the process of transcription of *CMK2* is not carried out. With less Cmk2 in the cell, inhibition of calmodulin does not occur, nor does the phosphorylation (and therefore inactivation) of the Pmr1 ATPase (Fig 15). From the activity of this cascade, we can infer that increased Far1 counteracts the effects of tamoxifen, explaining the increased growth rate in the *sam2Δ/sam2Δ* mutant cells. In fact, previous studies have shown that increased calmodulin suppresses tamoxifen activity [243].

**Fig 15.**
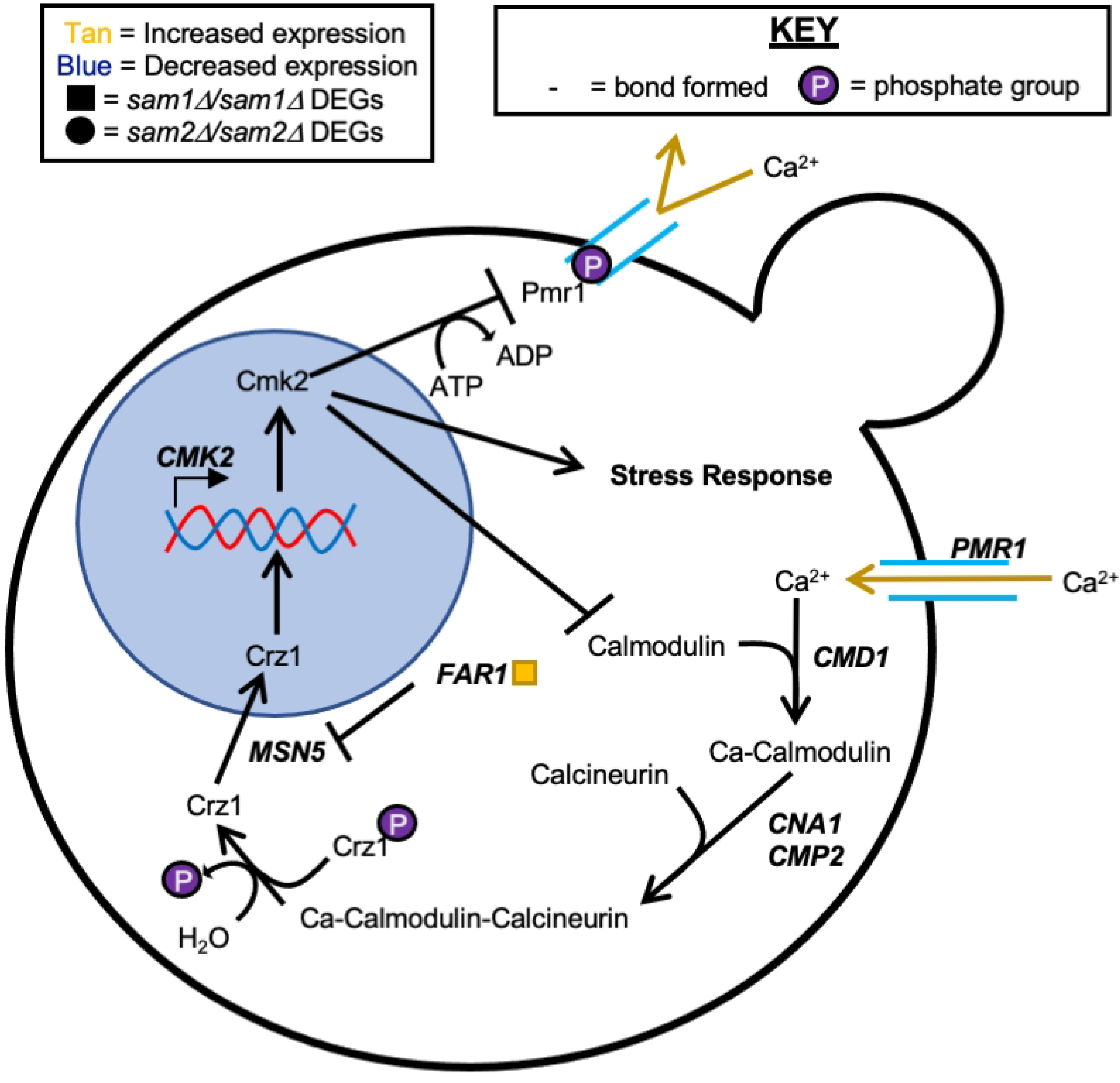
Calcineurin Signaling Cascade. Calcium ion signaling in *S. cerevisiae* begins with calcium import by membrane transporters. The calcium ion triggers a signaling cascade with calcineurin and calmodulin, which in response allows for cellular stress response. The gene names that encode the enzymes that can function in compound conversion in the pathway are given by their standard names. *sam2Δ/sam2Δ* DEGs are represented by red (increased expression) and green (decreased expression) squares. RNA-sequencing did not reveal any differentially expressed genes within this pathway between *sam1Δ/sam1Δ* and wildtype.

As well, previous work on tamoxifen resistance and sensitivity in yeast has been carried out in *Schizosaccharomyces pombe*. Two AdoMet-dependent methyltransferase genes, *EFM6* and *BUD23,* when deleted from *S. pombe,* result in tamoxifen resistance. *S. cerevisiae* homologs for these genes exist, with the same names, and which show significant decreases in expression in our *sam2Δ/sam2Δ* mutant strain [233]. Both encode methyltransferases involved in translation. *EFM6* showed a fold change in expression of 0.63 (p-value = 0.03995) while *BUD23* showed a fold change in expression of 0.36 (p-value = 0.00055). Neither *EFM6* nor *BUD23* exhibited any differential expression in the *sam1Δ/sam1Δ* cells (Supplementary Table 1). The direct mechanism for how decreased functionality of Efm6 and Bud23 lead to tamoxifen resistance has not been uncovered. What is known is that *EFM6* encodes for an AdoMet-dependent lysine methyltransferase, which modifies and activates translation elongation factor EF-1 alpha (eEF1A) by modifying the Lys-390 amino acid [57]. This enzyme is responsible for delivering aminoacyl-tRNAs to the ribosome during protein synthesis [247]. Furthermore, eEF1A is also responsible for binding to actin to assist with cytoskeleton organization and cellular shape and mobility [248]. *BUD23* encodes for a ribosome biogenesis factor and methyltransferase that methylates 18S rRNA, which is necessary for rRNA processing and export of the 40S ribosomal subunits from the nucleus [249]. As previously mentioned, *sam2Δ/sam2Δ* mutant cells have decreased AdoMet levels, which indicates that not only are the methyltransferases Efm6 and Bud23 decreased, but their methyl donor is as well. Because of this, we hypothesize that *sam2Δ/sam2Δ* mutant cells are reduced in their ability to carry out these processes via Efm6 and Bud23, contributing to the tamoxifen resistance of these cells. However, no known and/or published mechanism exists for why tamoxifen resistance was observed due to deletion of these genes in *S. pombe*, and while we see the same effect in *S. cerevisiae*, a known link has not previously been reported. Interestingly, Bud23 is known to interact with the rRNA methyltransferase Trm112, which may suggest that methylation further plays a role in ribosomal biogenesis [250]. Further, *TRM112* is significantly downregulated (fold change of 0.48, p-value=0.00075) in *sam2Δ/sam2Δ* mutant cells. As previously mentioned, Efm6 is also responsible for ribosome biogenesis, so we therefore hypothesize that tamoxifen may have some impacts on this process, and that downregulation of these genes somehow inhibits the toxicity of tamoxifen, though further research on this mechanism is warranted.

## DISCUSSION

Methylation is a frequently used regulation mechanism applied to practically all types of cellular macromolecules and used in a wide range of processes and pathways. S-Adenosylmethionine is the main methyl donor in all cell types, and cells therefore have a wide range of AdoMet-dependent methyltransferases to carry out these methylation reactions. Our previous research had identified loss of the two AdoMet synthetase encoding genes, *SAM1* and *SAM2*, as resulting in opposite impacts on AdoMet concentration and genome stability. Here we have presented further characterization of cellular changes due to loss of these two genes, through phenotypic profiling and RNA-sequencing. Of the 1,005 conditions tested via the Phenotypic Microarray we saw altered growth in ∼10% due to *sam1*-deficiency and ∼15% due to *sam2-*deficiency. The large number of conditions that result in altered growth speaks to the wide array of impacts that altering methyl donor abundance can impart, even when the conditions tested were not specifically selected as targeting known methyl involving pathways. Our RNA-sequencing experiment was able to capture expression from 7,127 genic regions, and we found 134 DEGs (1.88%) in our *sam1Δ/sam1Δ* strain and 876 DEGs (12.29%) in our *sam2Δ/sam2Δ* strain. This large percentage of altered gene expression due to *sam2*-deficiency again points to the broad impacts of the gene. Further, *sam2*-deficiency results in ∼50% more PM condition growth differences and approximately 6.5 times as many DEGs, demonstrating that its loss has greater cellular impacts, some of which are through decreased AdoMet concentration. Interestingly, we found 15 AdoMet-dependent methyltransferases downregulated, and 3 upregulated, in our *sam2Δ/sam2Δ* cells, as well as one upregulated and one downregulated non-AdoMet-dependent methyltransferase. This points to a mechanism whereby AdoMet concentration is involved in regulating their transcription, possibly via involvement with binding of activators or repressors or through altered chromatin modifications through reduced methyl donor availability for histone methylation.

The novel approach we have presented here, which combines data sets characterizing the broad changes resulting from loss of these genes, allowed us to more widely profile the variety of other impacts that were induced. By utilizing phenotypic profiling and gene expression profiling, we have provided insight that neither data set alone could provide. We have used the information from differential gene expression to show how the phenotypic changes are brought about, and have used the phenotypic profiling to show where the gene expression differences are large enough, individually or in combination, to cause altered cellular responses. Some of these changes likely contribute to the genome stability differences previously observed, while others provide insight on additional pathways disrupted by loss of one or both AdoMet-synthetase genes. Broadly, changes we have characterized are related to 1) AdoMet availability related to methyl cycle components as precursors in synthesis of other important molecules, 2) AdoMet availability relative to methylation reactions and methyltransferase functionality, and 3) impacted genes/pathways with no already defined relationship to methylation, the methyl cycle, or AdoMet itself. We have illustrated six stories to give multiple examples in each of these categories.

### Azoles

Azoles are synthetic organic compounds that have the ability to disrupt membrane integrity and ergosterol biosynthesis by targeting lanosterol 14α-demethylase, encoded by *ERG11* (Fig 5). Fluconazole is one of the most commonly prescribed medications to treat fungal and yeast infections as most yeasts, including *Saccharomyces cerevisiae*, treated with this azole drugs are normally susceptible. Previous studies have elucidated diverse mechanisms of azole resistance including, overexpression/deletion of genes involved in the ergosterol biosynthesis pathway or drug efflux pumps, as well as altered activity of the Set1 histone methyltransferase, or altered vacuolar sequestration [251–254]. We have identified a new mechanism of azole resistance; azole resistance in cells deficient in *sam2* occurs through the combined effects of decreased levels of AdoMet and the role the Erg6 methyltransferase plays in the alternate pathway.

The importance of the ergosterol biosynthesis pathway can be understood by recognizing the number of essential genes that are involved; deletion of any of 16 out of the 25 total genes result in cell death (Fig 5), due to lack of production of necessary metabolites [255–273]. While cells with mutations in any of the remaining 9 nonessential genes survive, they have been reported to have various defects such as; hypersensitivity to drugs and stress, decreased mating and transformation frequency, decreased tryptophan uptake, slowed growth, altered sterol levels, accumulation of aberrant sterols, lack of ergosterol, abnormal budding patterns, decreased fitness, altered gene expression and abnormal lipid molecules [37,77,274–291]. Although these defects have been shown to be correlated with lack of these nonessential genes, we don’t hypothesize the impacts to this pathway are linked to genome instability based on the observed phenotypes to date.

Our novel finding of impacting the ergosterol synthesis pathway through AdoMet and methyltransferase activity (not through a direct pathway mutation), to a large enough extent to result in increased sensitivity (via additional AdoMet) or increased resistance (via decreased AdoMet) has the potential to impact use of azole drugs moving forward. S-Adenosylmethionine is a common over-the-counter nutritional supplement and has also seen increased usage in treatment of a variety of ailments from liver disease to cancers to neurocognitive disorders in recent years [292–295]. Our studies suggest that addition of AdoMet to fluconazole/azole drug treatment might be effective in increasing sensitivity of other yeasts, like we have shown occurs in *S. cerevisiae*, and is worth further exploration.

### Cisplatin

Cisplatin is a strong platinum-based drug often used in chemotherapy [144]. It was introduced in 1845 then approved for medical use by the FDA in 1978 and is still widely used today in chemotherapeutic treatments. Cisplatin is most often used to treat cancers of the bladder, testicles, head/ neck, lungs, and ovaries [296]. Once in a cell, cisplatin causes damage to DNA [147]. Consequently, the DNA is unable to be replicated or transcribed, which may cause cell cycle arrest and apoptosis [147,148]. These attributes lead to its usage in cancer treatment with the goal to hinder the more actively dividing cancerous cells while minimizing damage to non-cancerous tissues. A balance must be sought in dosage to cause enough impact to cancerous cells that death ensues, but not be so overwhelming as to cause massive death to normal cells [297]. This leads to the potential for treatment to not kill all cancerous cells and the damage caused to their DNA to lead to novel mutations that might assist these cells in surviving further treatment.

As in *S. cerevisiae*, evidence in human cells shows that cisplatin uptake can occur through facilitated transporters such as the copper transporters [298]. Previous research has identified several multidrug exporters in human cells that exhibit upregulation in cells resistant to cisplatin [299]. We believe our observation that reduced AdoMet levels result in decreased expression of the *CTR1,* main copper transporter, as well as increased expression of drug efflux pumps, with an ultimate impact of reduced sensitivity to cisplatin, warrants further exploration as a cisplatin resistance mechanism in other organisms. Again, the availability of AdoMet as a nutritional supplement and in cancer therapeutics, would allow for exploration of its use as an adjuvant with cisplatin treatment. Further, our understanding that AdoMet availability in *S. cerevisiae* does not impact expression of DNA repair proteins or checkpoint genes, limits the worry that this might be a mechanism for resistance if supplemented with cisplatin treatment for cancers.

Regarding our interest in impacted pathways that might contribute to the genome stability changes observed in our mutant cells we do not think this condition has uncovered a linked process. While we were initially intrigued with altered growth patterns due to a DNA damaging agent, we saw resistance in our cells with more instability. These *sam2Δ/sam2Δ* mutants are less sensitive due to decreased import and increased export of cisplatin, not altered repair mechanisms we could see linked to instability.

### Glutathione

Glutathione (GSH) possesses anti-oxidative properties that are important in protecting cells from oxidative stress. In its reduced form, GSH is a reactant needed to neutralize reactive oxygen species (ROS), such as hydroxyls and superoxides [178]. ROS interact with proteins inducing a loss of protein function, cross-linking, and fragmentation causing increased levels of proteolysis. Additionally, ROS can interact directly with nucleic acids in DNA, which has been shown to lead to mutagenesis, carcinogenesis, and apoptosis, and ultimately genome instability. A known connection between GSH and the AdoMet cycle has been established through the transsulfuration pathway (Fig 10). The transsulfuration pathway plays a crucial role in both sulfur metabolism and redox regulation [99].

Sodium selenite and methyl viologen can both increase ROS levels and decrease the cells’ ability to combat ROS, while thiourea’s actions are concentrated on decreasing the cells’ ability to neutralize different ROS. All three of these conditions have the ability to increase genome instability through ROS. Sodium thiosulfate decreases ROS by readily donating electrons to the radical ROS species, inactivating them and rendering them harmless [176]. Our *sam2Δ/sam2Δ* mutants showed better growth in sodium thiosulfate/decreased ROS, while showing sensitivity to sodium selenite. The decreased expression of genes in the methyl cycle and in the synthesis of glutathione from homocysteine (and increased expression of those involved in transforming glutathione back to homocysteine) could point to decreased GSH to combat increased ROS. This pathway could directly be linked to the observed genome instability in these mutants and is under investigation by our group.

Interestingly we also saw sensitivity to conditions of increased ROS in our *sam1Δ/sam1Δ* cells. However, the growth patterns do not match what we saw in *sam2*-deficient cells, and the greatest sensitivity was seen in methyl viologen, where we could not observe a difference in growth in our *sam2Δ/sam2Δ* strain. This rules-out the possibility that the sensitivity mechanism might be the same in both mutants, and thus we do not hypothesize the mechanism in the *sam1*-deficient cells is linked to the GSH/methyl cycle but rather by a yet to be determined mechanism, and further research to discover the link is warranted.

### Arginine Metabolism

The arginine metabolism pathway includes the urea cycle and arginine biosynthesis, and further links to glutamate degradation, polyamine synthesis and the TCA cycle. Arginine metabolism is essential to a cell’s ability to respond to stress factors, regulate metabolic pathways, divide, and synthesize polyamines [191]. The most well understood connection to our *SAM* genes of interest is the involvement in polyamine synthesis which is also linked to the methyl cycle and AdoMet availability. The most common polyamines are putrescine, spermidine, and spermine, all of which are vital to genome stability and membrane stabilization. Ornithine serves as the direct link between the urea cycle/arginine biosynthesis and polyamine production (Fig 11). It is a precursor to putrescine, an essential precursor to the polyamines spermine and spermidine [192]. Synthesis of polyamines allows for increased DNA repair, stability, and resistance to genotoxic and oxidative damaging substances [193]. For example, evidence suggests that polyamines may promote the Rad51 genome repair complex by exposing damaged DNA to the repair system [192]. Furthermore, raised levels of polyamines in cells have been shown to increase rates of transcription of antioxidant enzymes, thereby lowering ROS [300]. Interestingly, despite an overwhelming increase in expression in most genes in the arginine biosynthesis pathway in our *sam2Δ/sam2Δ* mutants, close observation of individual reactions could point to a decrease in available ornithine. Not only is *CAR1* downregulated, responsible for direct conversion of L-arginine to ornithine, but the genes responsible for the reactions that convert ornithine to L-citrulline and further to L-arginino-succinate and L-arginine are all upregulated (*ARG3*, *ARG1*, and *ARG4* respectively). Decreased available ornithine, in tandem with the previously reported decrease in AdoMet in our *sam2Δ/sam2Δ* mutants, could point to decreased components for polyamine synthesis. This could be directly tied to the observed genome instability in these mutants, due to polyamine roles in increased DNA repair, stability and resistance to oxidative damage, and this pathway is under further investigation by our group.

### dNTPs

Ribonucleotide reductase (RNR) is the rate limiting enzyme required for *de novo* dNTP production. RNR is also responsible for monitoring and balancing dNTP pool size and is regulated via allosteric, localization, and transcriptional control. dNTP levels within the cell must be highly controlled in order to maintain genomic stability [204]. Each of the dNTPs, a small protein called Sml1, and ATP are the allosteric regulators of RNR [301,302]. Furthermore, RNR is able to detect dATP/ATP ratios via a feedback mechanism where dATP is an RNR inhibitor and ATP is an RNR activator [209]. Reduced dNTP pools have been shown to decrease DNA synthesis, thus decreasing the cell’s ability to repair DNA and perform recombination [303–305]. Research by [306] suggests that cells can sense dNTP levels and will arrest DNA strand elongation, via halting the replication fork, prior to the complete utilization of remaining dNTPs. Previous work has also demonstrated a mutator phenotype when dNTP levels exist at higher-than-normal amounts [301,305,307,308]; attributed to increased dNTPs speeding up S phase, increasing DNA Pol binding/extension from inaccurate primer-template pairing, and reduced proofreading efficiency [309–312].

5-fluorodeoxyuridine, 5-fluorocytosine, 6-azauracil, and hydroxylamine all inhibit DNA synthesis through altering deoxyribonucleoside triphosphate (dNTP) levels. dNTP pools are controlled by the *de novo* biosynthesis pathway and the salvage pathway. The pyrimidine salvage pathway allows for the catabolism of 2-deoxycytidine to form pyrimidine bases uracil, thymine, and cytosine. The *de novo biosynthesis of purines* pathway is not pictured in Fig 13, but also feeds in by providing the substrate PRPP, thus is still important in controlling dNTP balance. The salvage and *de novo* pathways interconnect with the folate cycle, and ultimately the methyl cycle, through shared intermediates. Because each of these pathways play a role in dNTP pool balance, they are likely involved in maintaining genomic stability as well. These pathways also connect to other important cellular pathways, such as arginine metabolism and nicotinamide and nicotinate metabolism (Fig 10), which provides insight to how dNTP pools relate to, and are affected by, several other processes within the cell.

While the resistance of our *sam2Δ/sam2Δ* mutant cells to many DNA damaging agents seems contrary to a link to the increased genome instability at first, we believe the contributing factor is likely in the dNTP pool regulation. We hypothesize the resistance to 5-fluorodeoxyuridine and 5-FC comes from upregulation of *CGI121* allowing for increase to production of dTTP, while 6-AU resistance is from increased expression of *SDT1* encoding pyrimidine nucleotidase which produces ribosides and also dephosphorylates the toxic form of 6-AU inactivating it. Further what we see at many steps in these interconnected pathways (Fig 13), is differentially expressed genes in the *sam2Δ/sam2Δ* mutants. This could potentially result in altered dNTP levels, and thus be directly linked to the observed genome instability in these mutants and is under further investigation by our group.

### Tamoxifen

Tamoxifen is a common chemotherapeutic drug, but also possess antifungal activity through inhibition of calmodulin binding to calcineurin [233,234]. Calcineurin and calmodulin play crucial roles in promoting calcium ion signaling pathways in the cell, which assist in regulation of enzymatic activity, metabolic functions, and stress response [235]. Previous work on tamoxifen resistance and sensitivity in yeast has been carried out in *Schizosaccharomyces pombe*. Two AdoMet-dependent methyltransferase genes, *EFM6* and *BUD23,* when deleted from *S. pombe,* result in tamoxifen resistance. *S. cerevisiae* homologs for these genes both encode methyltransferases involved in translation, and potentially involved in ribosome production and genome stability as well. The direct mechanism for how decreased functionality of Efm6 and Bud23 lead to tamoxifen resistance has not been uncovered, and no known and/or published mechanism exists for why tamoxifen resistance was observed due to deletion of these genes in *S. pombe*. While we see the same effect in *S. cerevisiae*, a known link has not previously been reported.

Both *EFM6* and *BUD23* also have identified homologs in humans [313,314]. Previous research has identified eukaryotic translation elongation factor 1-alpha 1 (eEF1A1) to be a novel inhibitor of the tumor-suppressor p53 gene, as well as causing increased resistance to chemotherapeutic drugs [315]. In *S. cerevisiae*, eEF1A1 is encoded by *EFM6*, and by the closest functional homolog, METTL21B in humans [316]. BUD23, the homolog of the same name in humans, has also been identified in influencing cellular proliferation and tumorigenesis [317]. Bud23 is methylated by Trm112 in *S. cerevisiae,* while Trmt112, the methyltransferase homolog, carries out this reaction in humans. *TRM112* and TRMT112 are AdoMet-dependent methyltransferases, which play a role in ribosome biogenesis and cell proliferation in both humans and *S. cerevisiae*, and have also been identified as being involved in tumor formation in humans [318]. The calcineurin signaling cascade is conserved across all eukaryotes [319]. In human cells, calcium ion signaling not only assists in enzymatic regulation, metabolic functioning, and stress response as seen in *S. cerevisiae*, but also aids in cellular proliferation, immune response, apoptosis, and cell differentiation [320]. Because the calcineurin signaling cascade is conserved, the actions that tamoxifen takes within a cancerous cell may be similar to the actions taken within *S. cerevisiae* cells. Previous studies have found that overactivation of the calcineurin signaling cascade promotes tumor growth by increased intracellular calcium ion, and therefore increased cellular proliferation [321]. One thing that should be brought to attention, however, is that traditionally, tamoxifen is used to treat breast cancer because it blocks estrogen receptors, which inhibits further cancerous growth and spread [322]. Further research has identified another mechanism of action for tamoxifen, in which the drug after long-term use has the capability to covalently bind to, and therefore alter the structure of, DNA and RNA, which could potentially cause further carcinogenesis [323,324]. We believe our observation that reduced AdoMet levels result in reduced sensitivity to tamoxifen, should be explored further as a tamoxifen resistance mechanism in other organisms, especially due to the recent increase of AdoMet usage in cancer treatments [325–327].

### Conclusions/Outlook

Our novel approach, combining Phenotypic Microarray data and RNA-sequencing data has elucidated several very promising avenues for further research. We have identified several pathways possessing altered gene expression in our mutants to explore in much greater detail to understand step-by-step how they might contribute to the observed genome instability in our mutants. We have also identified mechanisms of resistance to three commonly prescribed drugs due to lowered AdoMet and the resulting altered signaling in *S. cerevisiae*. The pathways these drugs act on have conserved components in *Homo sapiens*, and we suggest investigation of the use of AdoMet supplementation to combat resistance and potentially increase effectiveness of, or combat resistance to, these compounds. Further, the combination of phenotypic profiling and RNA-sequencing methodologies could be executed to learn vast amounts of information about all sorts of organisms and genes. We know our data sets contain additional stories that we plan to investigate, and those told here only scratch the surface of what we will be able to uncover.

## MATERIALS AND METHODS

### Strains

The *S. cerevisiae* strains used in this study are based on our W303 wildtype strain: MAT a/α, *leu2-3/leu2-3, his3-Δ200/his3-Δ200, trp1-Δ1/trp1-Δ1, lys2-801/LYS2, ura3-52/ura3-52, can1-100/CAN1, ade2-101/ade2-101, 2x [CF:(ura3::TRP1, SUP11, CEN4, D8B)].* The homozygous deletant strains for *SAM1* and *SAM2*, were created as described in [2]. Briefly, *SAM1* was knocked out with a KANMX cassette conferring resistance to G418: *sam1*::KANMX, and *SAM2* was knocked out with a *LEU2* cassette correcting the leucine auxotrophy: *sam2*::*LEU2*. Throughout this manuscript these strains are referenced by their *SAM1* and *SAM2* gene status: wildtype, *sam1Δ/sam1Δ,* and *sam2Δ/sam2Δ*.

### RNA Extraction and Quality Assessment

To prepare our yeast strains for RNA extraction, samples were grown in synthetic complete (SC) media altered to match their genotypic requirements (*sam1Δ/sam1Δ*: addition of G418, and *sam2Δ/sam2Δ*: removal of leucine). Samples were grown at 30°C overnight with agitation to mid-exponential phase (OD600 = 0.5-0.6). 1.5 mL of each culture was spun down at 14,000 rpm for one minute. The media was decanted and 750 µL of TRIzol was added. Samples were resuspended via pipetting and 250 µL of SIGMA acid-washed glass beads, 425-600 µm in size, were added. Samples were bead-beaten for 15 seconds, then rested on ice for 5 minutes, beaten again for 25 seconds, rested on ice for 5 minutes, then beaten for a final 15 seconds and placed back on ice. All remaining steps were conducted in a cold room at 4°C unless otherwise noted. 150 µL of chloroform was added to each sample, tubes were securely capped and vigorously shaken for 15 seconds. Samples were incubated for 2-3 minutes. Samples were then centrifuged at 12,000 × g for 15 minutes. The aqueous phase of each sample was transferred to a clean microcentrifuge tube and 375 µL of 100% isopropanol at -20°C was added. The samples were resuspended and incubated for 10 minutes before being centrifuged at 12,000 × g for 10 minutes. The supernatant was removed from the tubes and the remaining RNA pellets were broken apart and washed with 750 µL of 75% ethanol. The samples were vortexed briefly and centrifuged at 7,500 × g for five minutes. The supernatant was removed, and the pellets were allowed to air dry for 5-10 minutes at room temperature. Pellets were resuspended in 20 µL RNAase-free ddH_2_O and incubated at 60°C for 15 minutes before being stored at -80°C. RNA quantity and quality were assessed via Nanospec following the manufacturers guidelines. RNA quality was further assessed by running 1 µg of each sample on a 1% TAE gel with bleach at 100V for 40 minutes [328].

### RNA-Sequencing Library Preparation

Samples were sent to the University of Louisville Genomics Core Facility for RNA-Sequencing. The TruSeq Stranded mRNA LT Sample Prep Kit - Set A and Set B with poly-A enrichment was used to prepare a library for RNA-Seq. Briefly, polyadenylated RNA was purified, fragmented, and primed. First and second cDNA strands were synthesized, purified, and 3’ ends were adenylated. Then, the samples were barcoded with Illumina TruSeq adapters and the dsDNA fragments were enriched via PCR. The samples were purified, and quality was assessed using the Agilent DNA 1000 kit. The fragment size of the samples was about 300-350 bp in length. Sample quantity was assessed via a standard curve method from qPCR using DNA standards from the KAPA Library Quantitation Kit for Illumina Platforms. Next, the samples were normalized and pooled. Finally, the libraries were denatured and diluted in preparation for sequencing.

### RNA-Sequencing

An Illumina NextSeq 500, using NextSeq 500/550 1×75 cycle High Output Kit v2, was used to sequence the prepared library. Triplicates of three strains were analyzed: wildtype, *sam1Δ/sam1Δ* and *sam2Δ/sam2Δ.* Thirty-six single-end raw sequencing files (.fastq) were analyzed using the tuxedo suite pipeline. The raw data was condensed into one single-end .fastq file for each triplicate giving nine files representing each mutant strain [329]. Quality control (QC) was performed using FastQC (version 0.10.1) and it was determined that trimming the samples was unnecessary [330]. Tophat2 (version 2.0.13) was used to align sequences to the *S. cerevisiae* reference genome assembly (Saccharomyces_cerevisiae.R64-1-1.dna.toplevel.fa) [331]. The tuxedo suite of programs, including cufflinks-cuffdiff2 (version 2.2.1) were used to identify differentially expressed genes (DEGs) between each mutant strain and wildtype [332,333]. Cutoff limits for significance were as follows: p-value ≤ 0.05, 0.67 ≤ |FC| ≥ 1.5. A multiple comparison correction (MCC) was also applied, and q-value determined. These data can all be found in Supplementary Table 1. As we are interested in investigating pathways that might involve multiple DEGs, we are including discussion of all gene expression differences that meet the p-value ≤ 0.05, 0.67 ≤ |FC| ≥ 1.5 cutoffs, rather than the more stringent MCC q-values. In this manner we can visualize pathways where multiple p-value level significant DEGs are found, lending support to these not being false positive results.

### Phenotypic Microarray

Phenotypic Microarray plates were obtained from BiOLOG to test our mutant strains for phenotypic differences under a variety of conditions. Each well contains the necessary requirements for growth as well as different substrates or differing concentrations of the same substrate, testing a variety of conditions from metabolism to drug sensitivity. Substrate contents of each well are listed in Supplementary Table 5; concentrations are proprietary and confidential to BiOLOG. PM1 and PM2A MicroPlates test carbon sources, PM3B MicroPlate tests nitrogen sources, PM4A MicroPlate tests phosphorous and sulfur sources, PM5 MicroPlate tests nutrient supplements, PM6-8 MicroPlates test peptide nitrogen sources, PM9 MicroPlate tests osmolytes, PM10 tests pH, and PM21D, PM22D, PM23A, PM24C, and PM25D test for chemical sensitivities. All tests were conducted in duplicate for all three strains of interest.

Plates were inoculated per manufacturer instructions and the genotypes of our strains. Briefly, a stock solution for PM1-8 was prepared by adding 5 mL of His (2.4 mg/mL), Trp (4.8 mg/mL), Ura (2.4 mg/mL), and Leu (7.2 mg/mL) to 580 mL of ddH_2_O. A stock solution for PM9+ was prepared by adding 5 mL of His (2.4 mg/mL), Trp (4.8 mg/mL), Ura (2.4 mg/mL), and Leu (7.2 mg/mL) to 580ml SC-5 solution. The inoculating fluid for PM1-2 was prepared by adding 20 mL of IFY-0 (1.2x), 0.32mL of dye mix D (75x), and 3.18 mL of ddH_2_O. The inoculating fluid for PM3-8 was prepared by adding 60 mL of IFY-0 (1.2x), 0.96 mL of dye mix D (75x), 3 mL of D-glucose (24x), and 6.54 mL of ddH_2_O. The inoculating fluid for PM9+ was prepared by adding 70 mL of SC medium (1.2x), 0.84 mL of dye mix E (100x), and 7.91 mL of ddH_2_O.

Each yeast strain was streaked on separate BUY agar plates from BiOLOG and grown for 24 hours at 30°C then subcultured to a new BUY plate for an additional 24 hours. Cells from subculture plates were transferred into the previously prepared stock solutions to obtain uniform suspensions with a turbidity of 62% transmittance. 0.50 mL of the cell suspension was added to 23.5 mL of PM1,2 inoculating fluid, and 100 µL was added to each well of PM1 and PM2. 1.50 mL of the cell suspension was added to 70.50 mL of PM3-8 inoculating fluid, and 100 µL was added to each well of PM3-8. 1.75 mL of the cell suspension was added to 82.25 mL of PM9+ inoculating fluid, and 100 µL was added to each well of PM9-10, 21-25. A BioTek ELx800 microplate reader was used to obtain optical densities (OD) of the wells in each plate at 0 hours, 24 hours, 48 hours, and 72 hours. All plates were sealed with parafilm and incubated at 30°C with agitation between OD readings. Prior to each OD reading, the cells were resuspended via pipetting. OD readings for biological replicates at each time point were plotted to generate growth curves. Any growth curves where either the mutant strain or wildtype strain biological replicates were not consistent were deemed inconclusive. Significant PM growth curves that were used in this manuscript were depicted using GraphPad Prism version 9.5.1 for Windows, GraphPad Software. Graphs were generated using the third order polynomial (cubic) equation, and each graph was interpolated with a 95% confidence interval. Additional growth curves can be found in Supplementary Tables 6 and 7.

## ACKNOWLEDGEMENTS

Work was supported by grant 1R15GM109269-01A1 from the NIH, National Institute of General Medical Sciences, and an Institutional Development Award (IDeA) from P20GM103436 (Martha Bickford, PI). Sequencing and bioinformatics support for this work provided by University of Louisville Genomics & Bioinformatics Cores through National Institutes of Health (NIH) grant P30GM106396 (Donald Miller, PI). The contents of this work are solely the responsibility of the authors and do not represent the official views of the NIH or the National Institute for General Medical Sciences (NIGMS).

